# Assessing Reproducibility of High-throughput Experiments in the Case of Missing Data

**DOI:** 10.1101/2021.07.10.451851

**Authors:** Roopali Singh, Feipeng Zhang, Qunhua Li

## Abstract

High-throughput experiments are an essential part of modern biological and biomedical research. The outcomes of high-throughput biological experiments often have a lot of missing observations due to signals below detection levels. For example, most single-cell RNA-seq (scRNA-seq) protocols experience high levels of dropout due to the small amount of starting material, leading to a majority of reported expression levels being zero. Though missing data contain information about reproducibility, they are often excluded in the reproducibility assessment, potentially generating misleading assessments.

In this paper, we develop a regression model to assess how the reproducibility of high-throughput experiments is affected by the choices of operational factors (e.g., platform or sequencing depth) when a large number of measurements are missing. Using a latent variable approach, we extend correspondence curve regression (CCR), a recently proposed method for assessing the effects of operational factors to reproducibility, to incorporate missing values. Using simulations, we show that our method is more accurate in detecting differences in reproducibility than existing measures of reproducibility. We illustrate the usefulness of our method using a single-cell RNA-seq dataset collected on HCT116 cells. We compare the reproducibility of different library preparation platforms and study the effect of sequencing depth on reproducibility, thereby determining the cost-effective sequencing depth that is required to achieve sufficient reproducibility.

## A Introduction

High-throughput technologies are an essential part of modern biological research. Yet, outputs from high-throughput experiments are quite noisy due to many sources of variation in the experimental and analytic pipelines (referred to as workflows thereafter) that generate the data. The reproducibility of outcomes from a high-throughput workflow across replicated experiments provides important information for establishing confidence in measurements and evaluating the performance of the workflow.

The performance of a high-throughput workflow can be affected by many operational factors, such as experimental platforms or protocols, parameter settings in experimental procedures and labs conducting the experiments. Understanding how these factors affect the reproducibility of the outcome is crucial for identifying potential sources of irreproducibility and designing workflows that produce reliable results.

The primary output from a high-throughput workflow typically consists of a list of candidates (e.g., genes) that are evaluated by the workflow and their scores or measurements (e.g., gene expression) assigned by the workflow. One way to assess the reproducibility of a workflow is to compute the Pearson or Spearman correlation between the scores obtained on two replicate samples from the workflow for all candidates. Another way is to profile how consistently candidates are ranked and selected in replicate experiments across a sequence of selection thresholds. Methods based on this sequential approach include the correspondence at the top (CAT) plot [Irizarry et al., 2005], the correspondence curve [Li et al., 2011], the Irreproducible Discovery Rate (IDR) [Li et al., 2011], the Maximum Rank Reproducibility (MaRR) [Philtron et al., 2018] and the correspondence curve regression (CCR) method [Li and Zhang, 2018]. Despite differences, both types of approaches assume that the same set of candidates are observed across all replicate samples. Though this assumption holds for some high-throughput experiments, it is very common that some candidates are not observed in all replicates due to underdetection. For example, most single-cell RNA-seq protocols experience high levels of dropouts, where a gene is observed at a low or moderate expression level in one cell but is not detected in another cell of the same cell type [Kharchenko et al., 2014]. These dropouts occur due to the low amounts of mRNA in individual cells and inefficient mRNA capture, as well as the stochasticity of mRNA expression, making a vast majority of genes report zero expression levels [Qiu, 2020, **?**].

Because the aforementioned methods do not take missing values into account in the reproducibility assessment, a common practice for dealing with missing values in reproducibility assessments is to consider only the candidates with non-missing measurements, even though missing values do contribute useful information about reproducibility [Wu et al., 2013]. However, when there are a large number of zeros, this approach can lead to problematic results. For example, if only a small proportion of measurements are non-zero on all replicates and agree well across replicates, but the rest are all observed only on a single replicate, then ignoring zeros can lead to a seemingly high reproducibility despite the large amount of discordance in many candidates. Consider a study of HCT116 cells [Wu et al., 2013] where single-cell RNA-seq libraries are made using TransPlex Kit and SMARTer Ultra Low RNA Kit and whether or not including zeros in the data in the calculation of correlation will lead to substantially different conclusions about the reproducibility of these platforms. For example, when transcripts with zero counts are included (24933 transcripts), the Spearman correlation of the gene expression levels between a pair of cells is lower for TransPlex (0.648) than for SMARTer (0.734), whereas it is the opposite when only transcripts that are expressed in both cells are included (TransPlex: 0.501 for 8859 non-zero transcripts vs SMARTer: 0.460 for 6292 non-zero transcripts). This seems to imply that TransPlex is less reproducible than SMARTer if zeros are included, but the opposite if zeros are excluded. However, if the Pearson correlation is used as the reproducibility criterion, it suggests that TransPlex is more reproducible than SMARTer regardless of the inclusion of zeros (Supplementary Table C6). This inconsistency makes it difficult to make informative comparisons of different workflows about their reproducibility. The same issue also arises when profiling the consistency of candidates’ ranking, as different ways to handle missing values can lead to different results. A principled approach that accounts for missing values in reproducibility assessment is desirable.

In this work, we assess how the reproducibility of high-throughput experiments is affected by the choice of operational factors when a large number of measurements are missing due to under detection. Our approach is built on the correspondence curve regression (CCR) model, which assesses how the probability that a candidate consistently passes selection thresholds in different replicates is affected by operational factors in high-throughput experiments. By evaluating this probability at a series of rank-based selection thresholds through a cumulative link model, CCR summarizes the effects of operational factors on the reproducibility of the workflow across candidates at different significance levels as regression coefficients, allowing the effects on reproducibility to be assessed in a concise and interpretable manner. Our method extends this model by incorporating candidates with unobserved measurements through a latent variable approach. In doing so, this extension allows missing data to be properly accounted for in the assessment of the impact of operational factors on reproducibility.

This article is organized as follows. In Section 2, we first discuss the setup of the problem and provide a brief overview of the correspondence curve regression. Section 3 presents our missing data framework and the estimation procedure. In Section 4, we use simulation studies to evaluate the performance of our method. In Section 5, we apply our method to the dataset from a single-cell RNA-seq study. Section 6 concludes the paper.

## B Method

We consider the output from high-throughput experiments generated from *S* workflows (*S* ≥ 2). All the workflows measure the same underlying biological process, but they differ in certain operational factors such as experimental protocols, measurement platforms, or experimental parameters. We denote the vector of operational factors for the workflow *s* as **x**^**s**^. Table B1 shows how a dataset in such a setting would look.

**Table B1:** Output from *S* different workflows with 2 replicates each.

|  | Workflow 1 |  | Workflow 2 |  | ... | Workflow S |  |
| --- | --- | --- | --- | --- | --- | --- | --- |
|  | Replicate 1 | Replicate 2 | Replicate 1 | Replicate 2 |  | Replicate 1 | Replicate 2 |
| Gene 1 | $y_{11}^1$ | $y_{12}^1$ | $y_{11}^2$ | $y_{12}^2$ | ⋯ | $y_{11}^S$ | $y_{12}^S$ |
| Gene 2 | $y_{21}^1$ | $y_{22}^1$ | $y_{21}^2$ | $y_{22}^2$ | ⋯ | $y_{21}^S$ | $y_{22}^S$ |
| Gene 3 | $y_{31}^1$ | $y_{32}^1$ | $y_{31}^2$ | $y_{32}^2$ | ⋯ | $y_{31}^S$ | $y_{32}^S$ |
| ⋮ | ⋮ | ⋮ | ⋮ | ⋮ | ⋮ | ⋮ | ⋮ |
| Gene $i$ | $y_{i1}^1$ | NA | NA | $y_{i2}^2$ | ⋯ | $y_{i1}^S$ | $y_{i2}^S$ |
| ⋮ | ⋮ | ⋮ | ⋮ | ⋮ | ⋮ | ⋮ | ⋮ |
| Gene $j$ | NA | $y_{j2}^1$ | NA | $y_{j2}^2$ | ⋯ | $y_{j1}^S$ | NA |
| ⋮ | ⋮ | ⋮ | ⋮ | ⋮ | ⋮ | ⋮ | ⋮ |

For each workflow, few replicates are available, such that the reproducibility of the findings identified in the workflow can be assessed across replicates. For each replicate, the output consists of a list of candidates, such as genes, and their significance scores, which are assigned by the workflow to indicate the strength of evidence for a candidate to be a true signal. The scores can be original measurements or statistics derived from them (e.g., p-value), and we use these scores as our data. Because missing data in high-throughput studies are often created due to underdetection, we assume that the unobserved candidates receive a score lower than all the observed candidates. If a candidate is observed in at least one replicate, then it is termed as a partially observed candidate; if it is under the detection limit in all replicates, then it is termed as a missing candidate. Because missing candidates are completely unobserved, the total number of missing candidates is unknown. Without loss of generality, we assume that a score of large value indicates strong evidence and receives a low value in its rank, i.e. the most significant candidate receives rank one. The case of the reversed order, where the least significant candidate receives rank one, is also discussed in Section 3.3.

We will first consider the case that each workflow has two replicates, as this is the case that most existing reproducibility assessment methods are designed for, then extend it to the case of more replicates in Section 3.4. Let 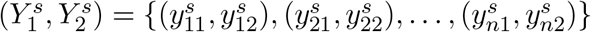 be the significance scores of a sample of *n* candidates on two replicates, assigned by the workflow *s*. We suppose that the scores 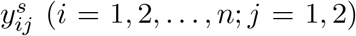, are a sample of the random variable 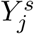 from an unknown distribution 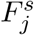. For notational simplicity, we omit the superscript *s* while describing a single workflow.

### B.1 Correspondence curve regression

Correspondence curve regression (CCR) is a cumulative link regression model that evaluates how covariates affect the reproducibility of the high-throughput experiments [Li and Zhang, 2018]. In such experiments, top candidates are often the targets for downstream analyses; hence, the consistency in the top-ranked candidates is important for reproducibility assessment. One challenge in this assessment is that the assessment results would be confounded with the threshold that is used to determine the cutoff for top ranks. CCR addresses this challenge by measuring the reproducibility at a series of thresholds. Specifically, it models the reproducibility at a given percentage threshold *t* as the probability that a candidate passes the specific threshold on both replicates,

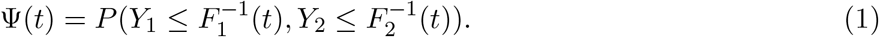

It then uses (1) as the response variable and models its relationship with operational factors at a series of thresholds using a cumulative regression framework. In doing so, the covariate effects on the response are reflected in the regression coefficients without being linked to any threshold, like cumulative regression models in other contexts. Given a set of pre-specified thresholds, 𝒯 = {*t*|0 < *t*_1_ <… < *t*_*M*_ ≤ 1}, the baseline model is defined as,

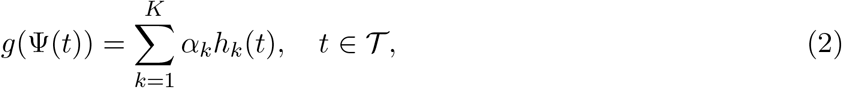

where (*h*_1_, …, *h*_*K*_) are pre-specified functions that describe the structure of the dependence between the replicates and (*α*_1_, …, *α*_*K*_) are unknown parameters that reflect the level of the dependence, i.e. the reproducibility of the workflow. The value of *K* and the functional forms of *h*_*k*_ and *g* can be determined based on the empirical data by an algebraic connection between (2) and Archimedean copula models [Li and Zhang, 2018] (introduced below). Now, let **x** be a vector of *d* covariates corresponding to operational factors for a workflow. For categorical variables, **x** will be the associated vector of dummy variables. The model with covariates is defined as,

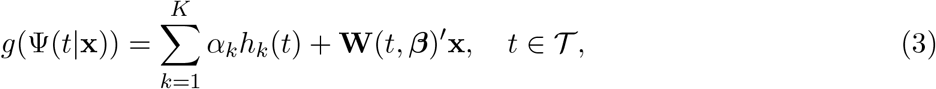

where **W**(*t*, ***β***)′**x** is a linear predictor characterizing the effect of the covariates **x** on reproducibility and ***β*** = (*β*_11_, …*β*_1*K*_, …, *β*_*d*1_, …*β*_*dK*_)^*T*^ are unknown coefficients to be estimated. Here, **W**(*t*, ***β***) = (*W*_1_(*t*, ***β***_1_), …, *W*_*d*_(*t*, ***β***_*d*_))^*T*^ with 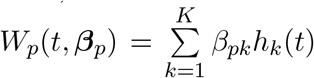, and *β*_*pk*_ measures the covariate effect on *h*_*k*_(*t*) due to *x*_*p*_ for *p* = 1, …, *d*.

To determine the functional form of (2), *Li and Zhang* [Li and Zhang, 2018] noticed that the response variable in (2) is in fact the diagonal section of a bivariate copula, i.e. Ψ(*t*) = *C*(*t, t*). Here, *C* is the bivariate cumulative distribution function of uniform random variables *T*_1_, *T*_2_ and *C*(*t*_1_, *t*_2_) = *P*(*T*_1_ ≤ *t*_1_, *T*_2_ ≤ *t*_2_). They showed that *g, α*_*k*_ and *h*_*k*_ in (2) take a canonical functional form if the dependence structure between *Y*_1_ and *Y*_2_ follows the dependence described by a certain Archimedean copula. Archimedean copulas are a parametric class of copulas that are used to model the dependence of multiple random variables. They take the form,

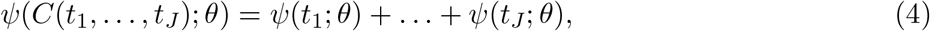

where *C*(*t*_1_, …, *t*_*J*_) = *P*(*T*_1_ ≤ *t*_1_, …, *T*_*J*_ ≤ *t*_*J*_), *θ* is an association parameter describing the strength of the dependence between uniform random variables *T*_1_, …, *T*_*J*_ and *ψ* is a generator function specific to each Archimedean copula. Different Archimedean copulas can describe different dependence structures. Thus, the functional form of (2) can be determined based on the copula that fits the empirical dependence structure of (*Y*_1_, *Y*_2_). To select the proper copula, one can plot the rank scatterplot of (*Y*_1_, *Y*_2_), which describes the empirical dependence structure of (*Y*_1_, *Y*_2_) and identify an Archimedean copula whose shape and tail behavior are similar to what is shown in the rank scatterplot. Figure 1 shows the rank scatterplots of data from two 1-parameter Archimedean copulas, the Gumbel-Hougaard copula and the Nelsen 4.2.12 copula. These two copulas reasonably capture the dependence structure of scores from replicated high-throughput experiments and can be represented in the form (2). See Table B2 for their respective functional forms. These two copulas differ in their shape and tail behavior. The Gumbel-Hougaard copula has higher concordance at the upper tail, whereas the Nelsen 4.2.12 copula has higher concordance at the lower tail. Hence, the Gumbel-Hougaard copula is suitable for the case when top candidates have large signals, and the Nelsen 4.2.12 is suitable for the case when top candidates have small signals, since top candidates usually have higher concordance across replicates than lowly ranked candidates. Consequently, the missing values due to underdetection tend to appear at the lower tail of the Gumbel-Hougaard copula and the upper tail of the Nelsen 4.2.12 copula, respectively, because missing values usually occur among the weaker signals.

**Figure B1:**
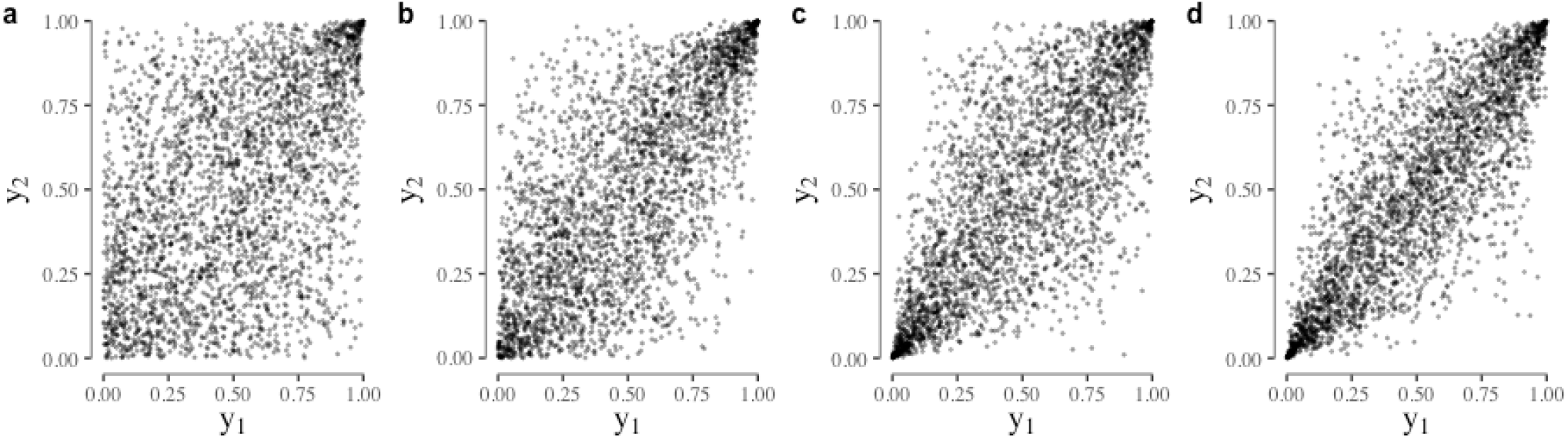
Rank scatterplots of data from the Gumbel-Hougaard copula and the Nelsen 4.2.12 copula with different association parameter *θ*. (a-b) Gumbel-Hougaard copula with association parameter *θ* = 1.5 and 2, respectively. (c-d) Nelsen 4.2.12 copula with *θ* = 1.5 and 2, respectively.

**Table B2:** Copula function *C*(·, ·; *θ*) and the corresponding canonical choices of *K, g* and *h*_*k*_(*t*) for the Gumbel-Hougaard copula and the Nelsen 4.2.12 copula, where *θ* is the association parameter for the copula models and *α*’s are the coefficients in (3).

| Copula name | $C(u, v; \theta)$ | $K$ | $g(\Psi(t))$ | $h_k(t)$ | $\alpha_k$ |
| --- | --- | --- | --- | --- | --- |
| Gumbel-Hougaard | $\exp(-[(-\ln u)^\theta + (-\ln v)^\theta]^{1/\theta})$ | 1 | $\log \Psi(t)$ | $h_1(t) = \log(t)$ | $\alpha_1 = 2^{1/\theta}$ |
| Nelsen 4.2.12 | $(1 + [(u^{-1} - 1)^\theta + (v^{-1} - 1)^\theta]^{1/\theta})^{-1}$ | 2 | $\log \frac{\Psi(t)}{1 - \Psi(t)}$ | $h_1(t) = 1, h_2(t) = \log \frac{t}{1-t}$ | $\alpha_1 = -\frac{\log 2}{\theta}, \alpha_2 = 1$ |

The CCR method has been shown to have good accuracy and power to estimate the difference in reproducibility between different workflows. Hence, we extend it to incorporate missing data.

## C Correspondence Curve Regression with Missing Data

### C.1 Missing data framework for upper tail truncation

As mentioned before, some candidates may be unobserved when signals are below the detection level. Consider the case when a small value of *y*_*ij*_ indicates strong evidence of a true signal (e.g., p-value) and receives a low rank. Then, a candidate *i* is observed in replicate *j*, if its score *y*_*ij*_ satisfies 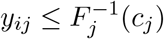. Here, *c*_*j*_ is unknown and needs to be estimated (see Section 3.2). Denote the corresponding observed score as 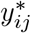, then we can write

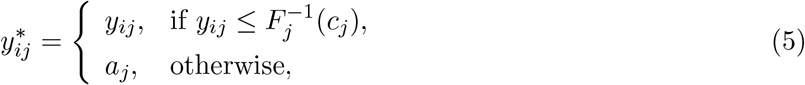

where *a*_*j*_ is a constant that satisfies 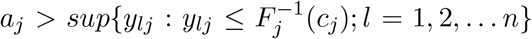 for replicate *j*. For example, if *y*_*ij*_ is a p-value and *c*_*j*_ < 1, then *a*_*j*_ can be 1. According to the missing status, candidates can be classified into three groups: the ones observed on both replicates, the ones observed on only one replicate but not the other, and the ones that are not observed on either replicate. They will be referred to as completely observed, partially observed and completely missing, respectively. Without loss of generality, we assume *c*_1_ ≤ *c*_2_. The sample space of (*Y*_1_, *Y*_2_) can be partitioned by *c*_1_ and *c*_2_ into four categories, *C*11, *C*12, *C*21 and *C*22, as shown in Figure 2.

**Figure C2:**
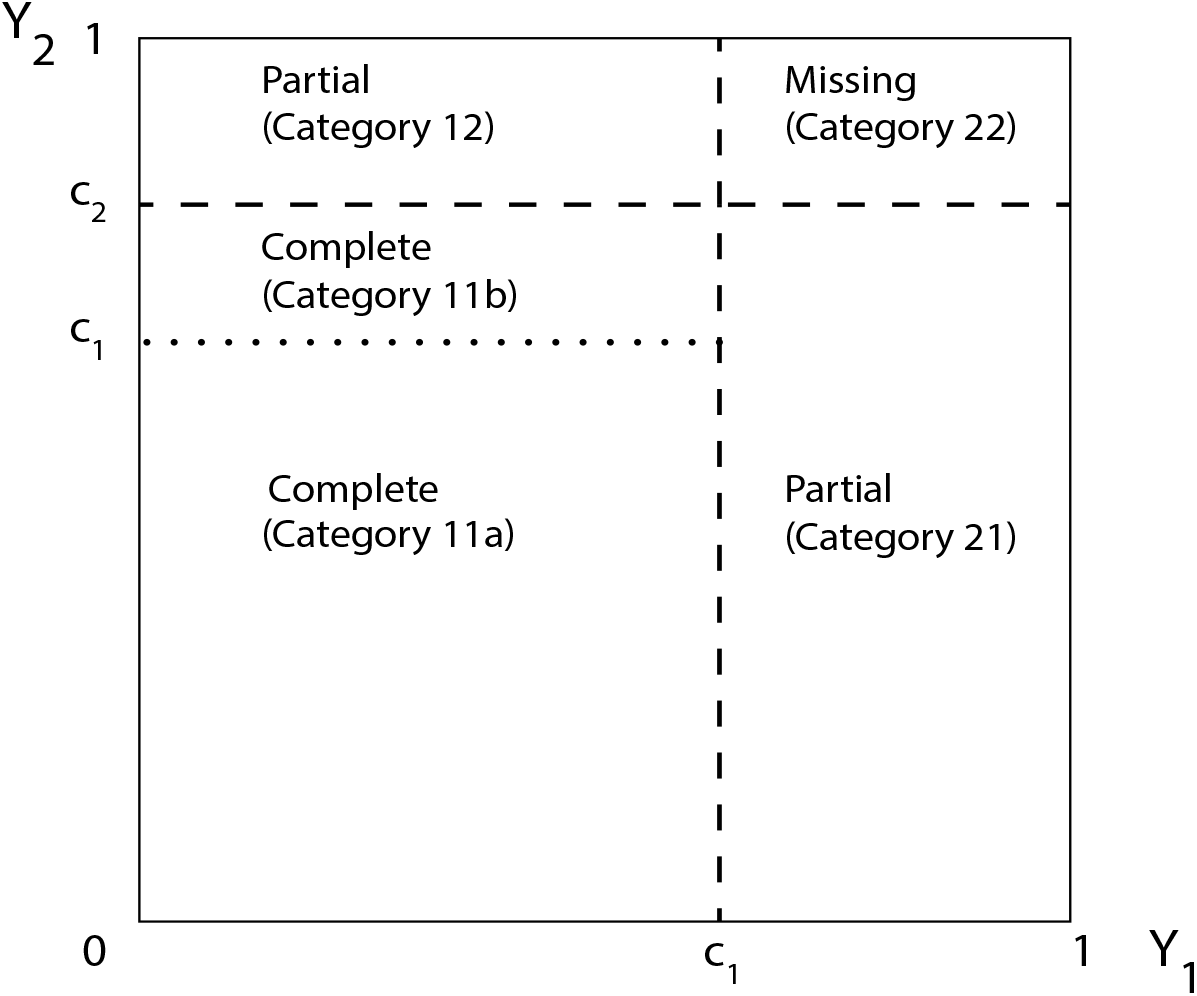
Partition of sample space according to missing status in the case of upper tail truncation.

Category 11: 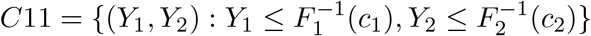, i.e. completely observed.

Category 12: 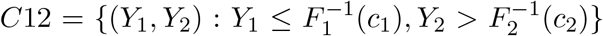, i.e. partially observed (observed on *Y*_1_ but missing on *Y*_2_).

Category 21: 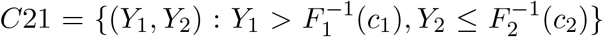, i.e. partially observed (observed on *Y*_2_ but missing on *Y*_1_).

Category 22: 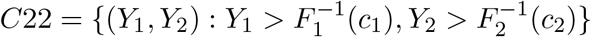, i.e. completely missing.

Denote the probabilities of falling into these categories as *P*_11_, *P*_12_, *P*_21_ and *P*_22_, respectively. Then the corresponding probabilities are as follows:

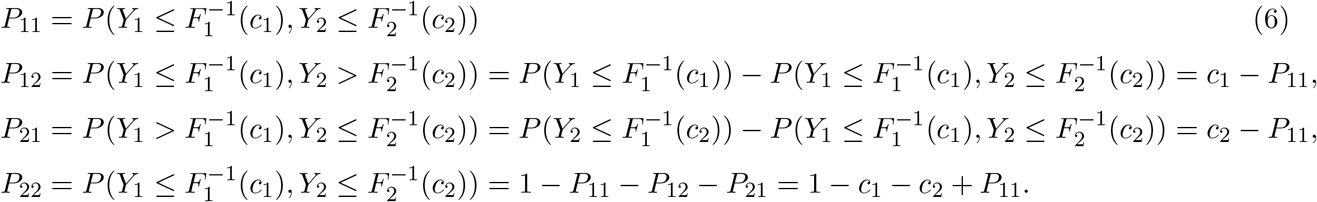

Denote the numbers of candidates in these categories as *n*_11_, *n*_12_, *n*_21_, and *n*_22_, respectively. Note that the occurrences of completely missing candidates are unobserved. Thus, if a category has completely missing candidates, the number of observations in this category is unknown. Hence, the values of *n*_11_, *n*_12_, *n*_21_ are known for a dataset, but *n*_22_ is unknown.

### C.2 Likelihood function with missing data due to upper tail truncation

We develop a maximum likelihood approach to estimate the parameters in this model. This approach is built upon the estimation strategy for the CCR model [Li and Zhang, 2018], which partitions observations into non-overlapping ordered categories and fits a multinomial likelihood. We extend this approach to include missing data information using the EM algorithm for truncated data [Dempster et al., 1977]. As we have shown in Section C.1, when there are missing data, the entire sample space can be partitioned into the category with completely observed candidates (Category 11) and the categories with partial and missing candidates (Category 12, 21, and 22). Therefore, we can conveniently extend the multinomial likelihood in the CCR model by incorporating partial and missing data as additional categories and treating the completely observed category in the similar way as in the CCR model as follows.

For notational simplicity, we first consider the likelihood for a single workflow. Denote the parameters to be estimated as *ϕ* = {***ω***, *c*_1_, *c*_2_}, where ***ω*** = (***α, β***) = (*α*_1_, …, *α*_*K*_, *β*_11_, …, *β*_*dK*_). To compute the multinomial likelihood, we first compute *P*_11_, which will be used to compute *P*_12_, *P*_21_ and *P*_22_ subsequently using (6). To proceed, we first partition *C*11 into two subcategories, *C*11^*a*^ and *C*11^*b*^, where 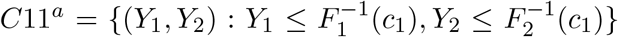 and 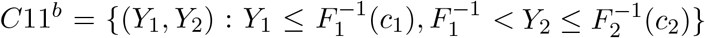. Denote the probability of these two categories as *P*_11*a*_ and *P*_11*b*_, respectively, and the corresponding counts as *n*_11*a*_ and *n*_11*b*_, respectively. Then

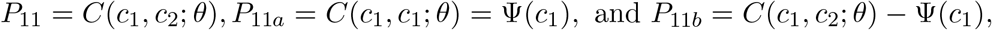

where Ψ(·) = *g*^−1^(Σ*α*_*k*_*h*_*k*_(·) + **W**(·, ***β***)′**x**) and *C*(·, ·) is the copula function corresponding to the regression model, with the copula association parameter *θ* obtained from the regression coefficient, (*α, β*), using the correspondence described in the CCR model (See Section B.1 and Table B2 last column, more details in the supplementary information A.1). For *C*11^*a*^, we partition it further into non-overlapping ordered categories using a series of cutoffs, as in *Li and Zhang* [Li and Zhang, 2018]. Specifically, considering the cutoffs, 0 < *t*_1_ < …< *t*_*M*_ = *c*_1_, the candidates from each workflow that fall in *C*11^*a*^ can be partitioned into M blocks, *C*111, …, *C*11*M*, such that each block consists of the candidates that are deemed reproducible at *t*_*m*_ but not at *t*_*m*−1_ (see Figure 2). The observations in block *m* can be represented as 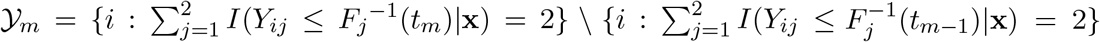, where *Y*_*ij*_ is the score of candidate *i* on replicate *j* and *F*_*j*_ is the empirical cumulative distribution function of *Y*_*j*_, *i* = 1, 2, …, *n*; *j* = 1, 2. Let *U*_*im*_ = *I*(*i* ∈ 𝒴_*m*_|**x**, *c*_1_, *c*_2_) be the binary indicator for candidate *i* to fall in the *m*^*th*^ block. That is, *U*_*im*_ = 1, if *y*_*ij*_ satisfies *t*_*m*−1_ < *F*_*j*_(*y*_*ij*_) ≤ *t*_*m*_ ∀*j*. Denote the number of observation in block *m* as *n*_11*m*_; *m* = 1, …, *M*, and the probability to fall into block *m* as *P*_11*m*_, then

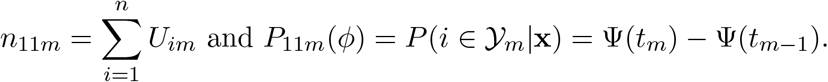

Unlike the case that all data points are observed in *Li and Zhang* [Li and Zhang, 2018], *F*_*j*_(·) is not observed directly due to truncation. Yet, we can compute *F*_*j*_(*y*) for any observed value *y* (i.e. 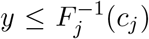) as 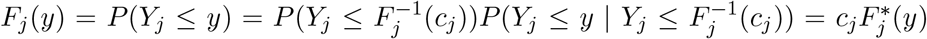, where 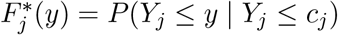 is the cumulative distribution of the truncated distribution. Because the observed values of *Y*_*j*_ are upper truncated at *c*_*j*_ for the *j*^*th*^ replicate, 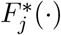 can be computed from the observed scores.

Following the EM algorithm for truncated data [Dempster et al., 1977], we augment the data to include the number of candidates in *C*22, *n*_22_, which is not observed due to truncation. Denote the total number of the candidates in categories *C*11, *C*12 and *C*21 as *n*_*obs*_ = *n*_11_ + *n*_12_ + *n*_21_, and the corresponding probability *P*(*ϕ*) = *P*_11_(*ϕ*) + *P*_12_(*ϕ*) + *P*_21_(*ϕ*). Then *n*_22_ ∼ Negative Binomial(*n*_*obs*_, *P*(*ϕ*)). The complete data likelihood is,

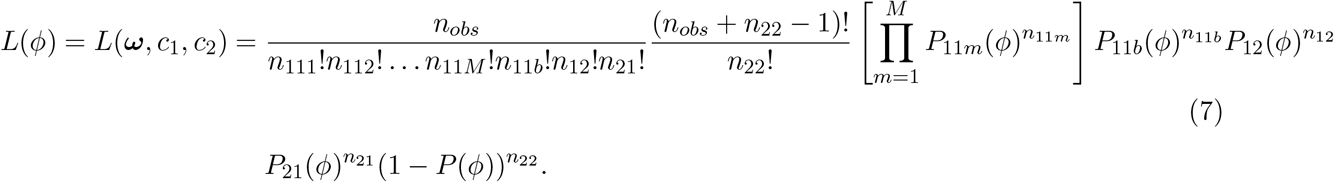

The detailed EM steps can be found in the supplementary information A.1.

When there are *S* workflows, the number of observed data from *S* workflows is denoted as 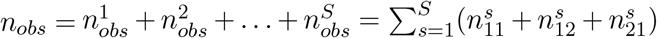 and the number of candidates in category *C*22 of the *s*^*th*^ workflow, 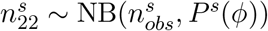. Then, the complete data likelihood for all workflows is,

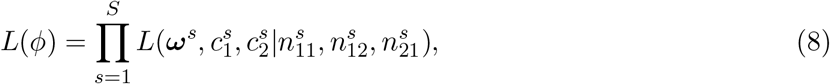

where 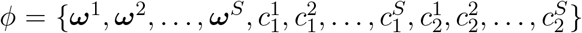, **Ω** = {***ω***^1^, ***ω***^2^, …, ***ω***^*S*^} is the vector of reproducibility parameters for *S* different workflows and 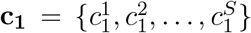 and 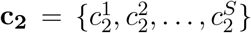 are vectors of truncation parameters for *S* different workflows. Supplementary information A.1 provides the details of the EM algorithm. We call our proposed method as the Correspondence Curve Regression with Missing Data (mdCCR).

### C.3 Inverted missing data framework for lower tail truncation

Sometimes, a large value of *y*_*ij*_ indicates strong evidence of a true signal and receives a high rank. In this case, a candidate *i* is observed in replicate *j*, if its score *y*_*ij*_ satisfies 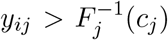, and the missing data would appear in the lower tail. Denote the corresponding observed score as 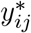, then we can write

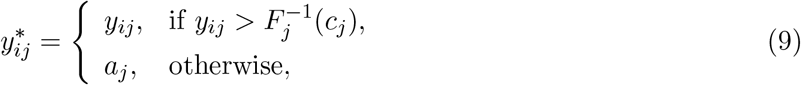

where *a*_*j*_ is a constant that satisfies 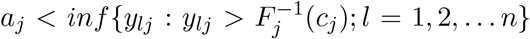 for replicate *j*. For example, if *y*_*ij*_ is a gene expression value and *c*_*j*_ *>* 0, then *a*_*j*_ can be 0. Without loss of generality, we assume *c*_1_ ≥ *c*_2_. The whole data set can be partitioned into four categories by *c*_1_ and *c*_2_ as shown in Figure 3. The categories and corresponding probabilities are as follows:

**Figure C3:**
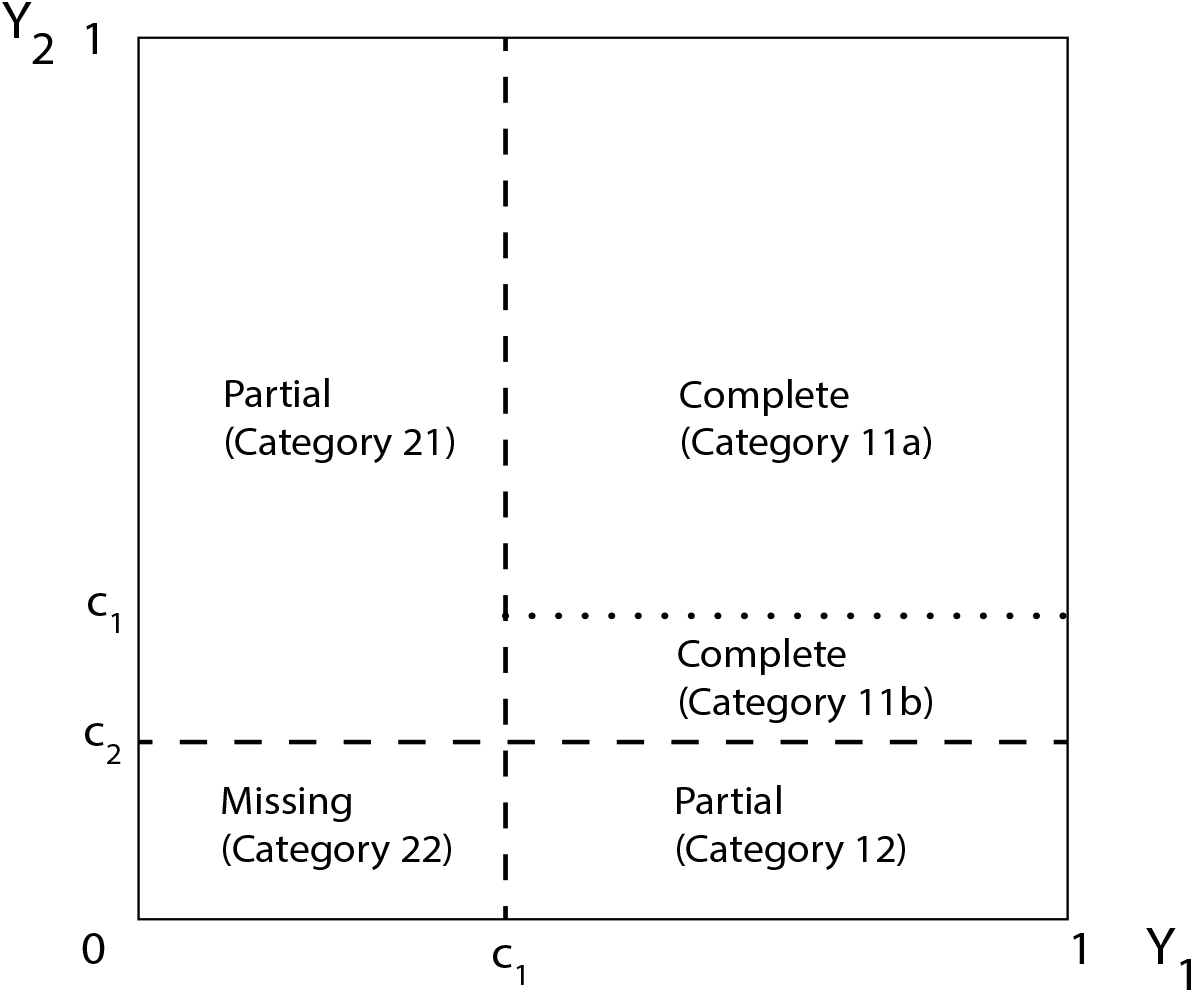
Partition of sample space in the inverted missing data framework (lower tail truncation) according to missing status.

Category 11: 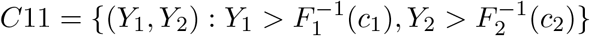 and *P*_11_ = 1 − *c*_1_ − *c*_2_+*C*(*c*_1_,*c*_2_)

Category 12: 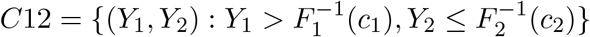 and *P*_12_ = 1 − *c*_1_ − *P*_11_

Category 21: 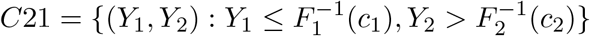 and *P*_21_ = 1 − *c*_2_ − *P*_11_

Category 22: 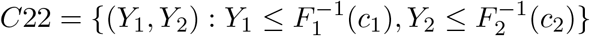 and *P*_22_ = *c*_1_ + *c*_2_ + *P*_11_ − 1,

where *C*(·, ·) is the copula function corresponding to the regression model, with the copula association parameter obtained from the regression coefficient using the correspondence described in the CCR model.

Denote the numbers of candidates in these categories as *n*_11_, *n*_12_, *n*_21_, and *n*_22_, respectively. The estimation for this case can be done in a way similar to the procedure in section C.2 with a few changes as follows. That is, *C*11 is partitioned into *C*11^*a*^ and *C*11^*b*^, where 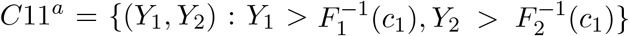 and 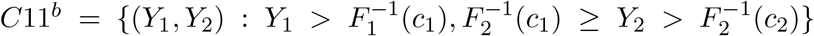. The corresponding probabilities are

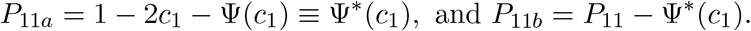

Then *C*11^*a*^ is further partitioned with the cutoffs of *c*_1_ = *t*_1_ < …< *t*_*M*_ = 1 to form blocks of 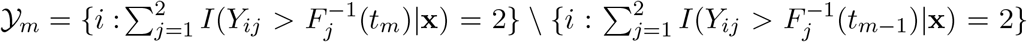. The probability of falling in category *C*11*m* is *P*_11*m*_(*ϕ*) = Ψ*(*t*_*m*−1_) − Ψ*(*t*_*m*_); *m* = 1, 2, …, *M*, where Ψ*(*t*_*m*_) = 1 − 2*t*_*m*_ + Ψ(*t*_*m*_).

Because the observed values of *Y*_*j*_ are lower truncated at *c*_*j*_ for the *j*^*th*^ replicate, for any observed *y*(i.e.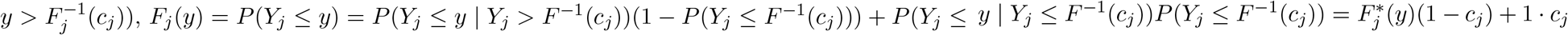, where 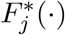 is the cumulative distribution of the truncated distribution and can be computed from the observed scores. The complete likelihood for *S* workflows can be computed using (7) and (8), similar to Section C.2.

### C.4 Framework for more than 2 replicates

Now we extend our model to the case that more than 2 replicate experiments are conducted by a workflow, a situation that is commonly encountered in some high-throughput settings. To assess the reproducibility across multiple replicate experiments, we leverage the information about concordance between pairs of replicates through a pairwise composite likelihood approach. Specifically, suppose that 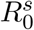 replicate experiments are conducted by the *s*^*th*^ workflow, then 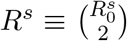 pairs of replicates can be formed. For each pair, the likelihood of the concordance between the replicates within the pair takes the form as described in the previous sections. Because each replicate is used in *R*^*s*^ − 1 pairs, the likelihood functions for pairwise concordance are not independent across some pairs. Hence, we combine information across pairs using a composite likelihood approach, which is a computationally efficient approach commonly used for approximating joint data models. Composite likelihood assumes independence across individual pairs. It has been shown that this approach produces consistent estimators, though the estimators exhibit some loss of efficiency [Varin et al., 2011]. If 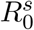 is moderate or large, one may use one replicate only once in one pair, then 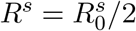. In this case, the independence assumption in the composite likelihood actually holds.

Specifically, for a given workflow *s*, we assume that all pairs of replicates share the same set of parameters 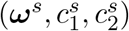, because all replicates are generated from the same workflow. Hence, the number of unobserved candidates for the *s*^*th*^ workflow from all replicates share the same distribution i.e., 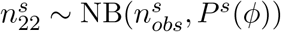, where 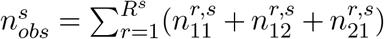. The complete composite likelihood is,

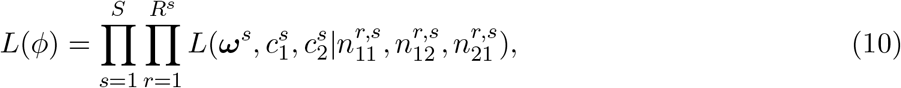

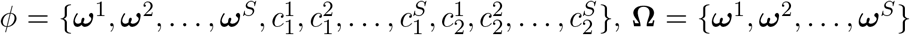 is the vector of reproducibility parameters and 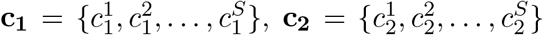 are vectors of truncation parameters for *S* different workflows. We estimate the parameters based on the observed data from all the replicates for *S* workflows using the maximum likelihood estimation via EM algorithm (see details in the supplementary information A.1).

## D Simulation studies

We use simulation studies to examine the performance of our method. In particular, we evaluate how missing data affect the accuracy of estimation, the type I error and the power for detecting differences in reproducibility in the settings resembling high-throughput experiments. To determine if the parameters are accurately estimated from the regression model, we generate (*Y*_1_, *Y*_2_) from the copulas discussed in Section B and then estimate the regression coefficient using the corresponding canonical functional choice. In this way, the accuracy of estimation can be determined by comparing the estimated parameters with the association parameter in the copula model.

We simulate the data from two copula models, the Nelsen 4.2.12 and Gumbel-Hougaard copula models. These models fit many high-throughput datasets and have been shown to be suitable for ChIP-seq and microarray datasets [Li and Zhang, 2018]. As mentioned earlier, the two copulas differ in their shape and tail behavior (Figure 1), as well as the location of the missing data in the structure. The Nelsen 4.2.12 copula has missing values at the upper tail and the Gumbel-Hougaard copula has missing values at the lower tail. Hence, the framework in Section C.1 fits the Nelsen 4.2.12 copula and the inverted framework in Section C.3 fits the Gumbel-Hougaard copula. After we generate (*Y*_1_, *Y*_2_) from the corresponding copula models, we create missing observations by removing observations under the detection threshold. That is, for each workflow, the missing observations are created by removing *Y*_1_ *> F* ^−1^(*c*_1_) and *Y*_2_ *> F* ^−1^(*c*_2_) and vice versa for the inverted case. In all simulations, *c*_2_ is kept equal to *c*_1_. We then estimate the regression coefficient using the corresponding regression model in Section D. As a comparison, we also perform the analysis using the CCR model, which excludes all the partially observed and missing data and only uses completely observed data.

### D.1 Accuracy of estimation and model fitting

To evaluate the accuracy of our estimation procedure, we simulate data from the baseline model (model with a single workflow i.e. (2)) with a single pair of replicates. We simulate from the Nelsen 4.2.12 (case 1) and the Gumbel-Hougaard copulas (case 2). For each copula, we choose the association parameter of the copula, *θ*, to be 2 and 3, as they reasonably reflect the level of dependence usually found in real data. We vary *c*_1_ such that 10%-40% of the data are missing in each replicate. For each case, we simulate 400 datasets, each of which consists of scores for *n* = 10000 candidates on a pair of replicates. We perform the estimation using our method and the CCR method with *M* = 30 equally spaced cutoff points in (0, 1) for both methods. The standard errors for the estimators in mdCCR are calculated using 400 bootstrap samples.

#### Case 1: *Nelsen 4*.*2*.*12 copula*

The baseline model is logit(Ψ(*t*)) = *α* + logit(*t*), where *α* = − log(2)*/θ, θ* ∈ [1, ∞). Here, *c*_1_ is chosen to be 0.6, 0.7, 0.8, 0.9, and the proportion of missing data increases as *c*_1_ decreases.

#### Case 2: *Gumbel-Hougaard copula*

The baseline model is log(Ψ(*t*)) = *α* log(*t*), where *α* = 2^1*/θ*^, *θ* ∈ [1, ∞). Here, *c*_1_ is chosen to be 0.1, 0.2, 0.3, 0.4, and the proportion of missing data increases as *c*_1_ increases.

We use mean integrated squared errors (MISE) as a metric to assess model fitting. MISE for Ψ(*t*) is the deviation between the empirical correspondence curve 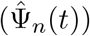 and the fitted curve 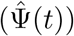, which is defined as,

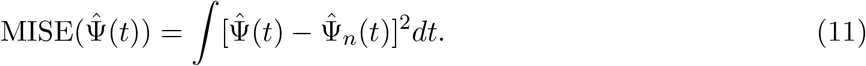

Here, we integrate over all the observed data to keep the comparison of the two methods on the same basis. For the inverted framework, we use MISE for Ψ*(*t*), i.e. 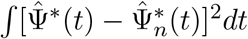.

As shown in Table D3, the estimate of *α* from CCR has a large deviation from the true value, whereas the estimate from our method is close to the true value at all levels of missing data. Our method also estimates the amount of missing data reasonably well. Furthermore, our method shows a much smaller MISE than CCR in all cases, indicating a better fit. The gain of our method over CCR increases with the increase of missing data.

**Table D3:**
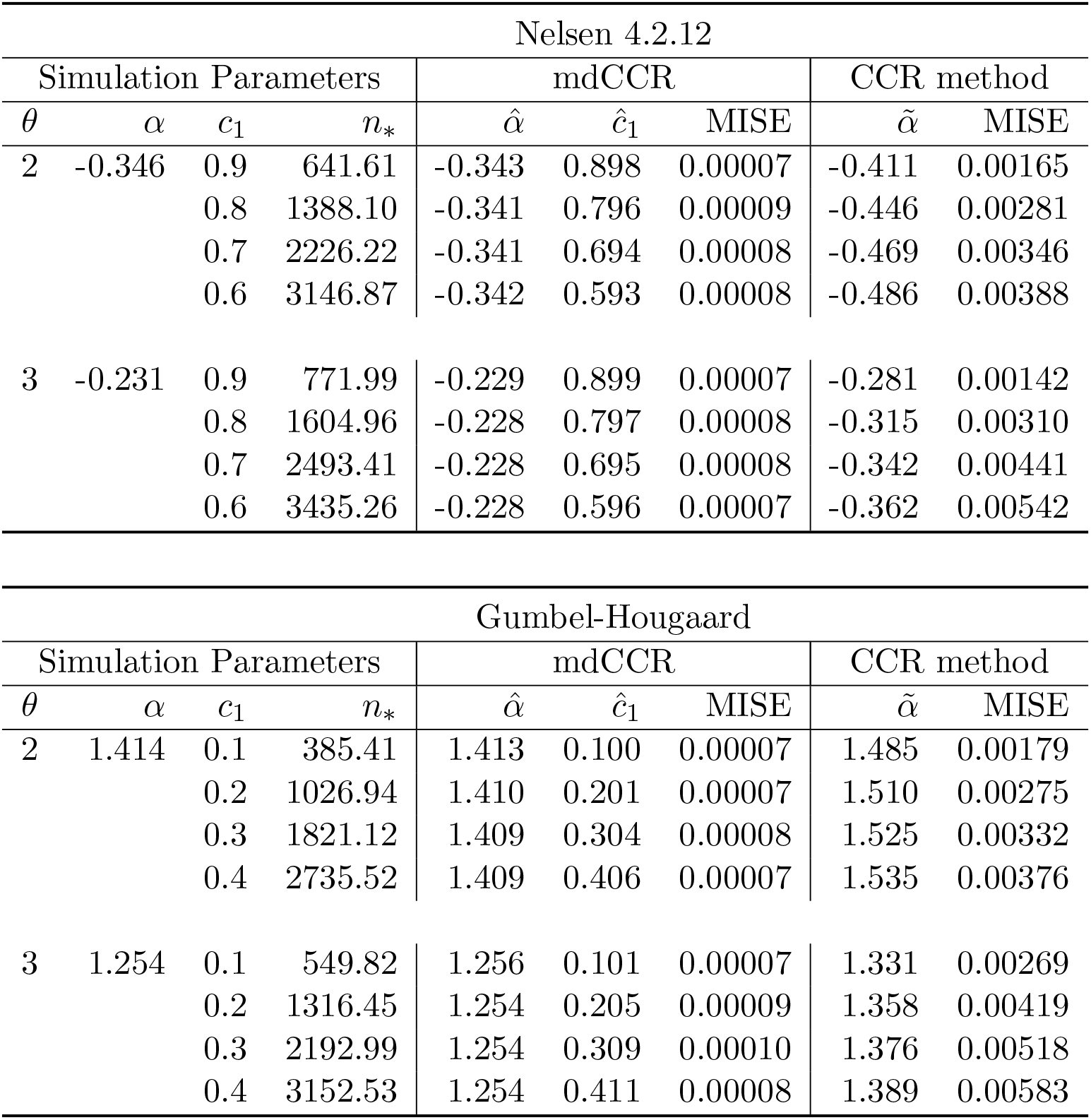
Comparison of estimates using mdCCR and CCR for baseline model of the Nelsen 4.2.12 and the Gumbel-Hougaard copula. 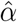 is the estimate using mdCCR, and 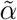 is the estimate using the CCR method.

| Nelsen 4.2.12 |  |  |  |  |  |  |  |  |
| --- | --- | --- | --- | --- | --- | --- | --- | --- |
| Simulation Parameters |  |  |  | mdCCR |  |  | CCR method |  |
| $\theta$ | $\alpha$ | $c_1$ | $n_*$ | $\hat{\alpha}$ | $\hat{c}_1$ | MISE | $\tilde{\alpha}$ | MISE |
| 2 | -0.346 | 0.9 | 641.61 | -0.343 | 0.898 | 0.00007 | -0.411 | 0.00165 |
|  |  | 0.8 | 1388.10 | -0.341 | 0.796 | 0.00009 | -0.446 | 0.00281 |
|  |  | 0.7 | 2226.22 | -0.341 | 0.694 | 0.00008 | -0.469 | 0.00346 |
|  |  | 0.6 | 3146.87 | -0.342 | 0.593 | 0.00008 | -0.486 | 0.00388 |
| 3 | -0.231 | 0.9 | 771.99 | -0.229 | 0.899 | 0.00007 | -0.281 | 0.00142 |
|  |  | 0.8 | 1604.96 | -0.228 | 0.797 | 0.00008 | -0.315 | 0.00310 |
|  |  | 0.7 | 2493.41 | -0.228 | 0.695 | 0.00008 | -0.342 | 0.00441 |
|  |  | 0.6 | 3435.26 | -0.228 | 0.596 | 0.00007 | -0.362 | 0.00542 |

| Gumbel-Hougaard |  |  |  |  |  |  |  |  |
| --- | --- | --- | --- | --- | --- | --- | --- | --- |
| Simulation Parameters |  |  |  | mdCCR |  |  | CCR method |  |
| $\theta$ | $\alpha$ | $c_1$ | $n_*$ | $\hat{\alpha}$ | $\hat{c}_1$ | MISE | $\tilde{\alpha}$ | MISE |
| 2 | 1.414 | 0.1 | 385.41 | 1.413 | 0.100 | 0.00007 | 1.485 | 0.00179 |
|  |  | 0.2 | 1026.94 | 1.410 | 0.201 | 0.00007 | 1.510 | 0.00275 |
|  |  | 0.3 | 1821.12 | 1.409 | 0.304 | 0.00008 | 1.525 | 0.00332 |
|  |  | 0.4 | 2735.52 | 1.409 | 0.406 | 0.00007 | 1.535 | 0.00376 |
| 3 | 1.254 | 0.1 | 549.82 | 1.256 | 0.101 | 0.00007 | 1.331 | 0.00269 |
|  |  | 0.2 | 1316.45 | 1.254 | 0.205 | 0.00009 | 1.358 | 0.00419 |
|  |  | 0.3 | 2192.99 | 1.254 | 0.309 | 0.00010 | 1.376 | 0.00518 |
|  |  | 0.4 | 3152.53 | 1.254 | 0.411 | 0.00008 | 1.389 | 0.00583 |

### D.2 Power and type I error for detecting differences in reproducibility

Next, we assess the power and type I error for detecting the difference in reproducibility. Here, two workflows are considered, so *x* takes value 0 or 1. Each workflow has two replicates. We define *α*_0_ and *α*_1_ as the coefficients related to *x* = 0 and *x* = 1, respectively. Similarly, *θ*_0_ and *θ*_1_ are the association parameters related to *x* = 0 and *x* = 1, respectively. We again simulate data from the Nelsen 4.2.12 and the Gumbel-Hougaard copulas with varying proportions of missing data. The corresponding regression model (3) and parameters for each case are summarized below.

#### Case 1: *Nelsen 4*.*2*.*12 copula*

The covariate term is **W**(*t, β*) = *α*_1_ + logit(*t*) − (*α*_0_ + logit(*t*)) ≡ *β* and so the model is logit(Ψ(*t*)) = *α*_0_ + *βx* + logit(*t*). *α*_*x*_ = − log(2)*/θ*_*x*_, where *θ*_*x*_ ∈ [1, ∞) for *x* = 0, 1 and *β* = *α*_1_ − *α*_0_ = − log(2)*/θ*_1_ − (− log(2)*/θ*_0_). This model indicates that the probability that a candidate is reproducibly ranked among top 100*t*% in workflow *x* = 1 is exp(*β*) times of that in *x* = 0. Hence, a positive value of *β* indicates that workflow *x* = 1 is more reproducible than workflow *x* = 0. Here, *c*_1_ is chosen to be 0.6, 0.7, 0.8, 0.9, 1, where a smaller *c*_1_ indicates a larger proportion of missing data and *c*_1_ = 1 represents the scenario that all the data are completely observed.

#### Case 2: *Gumbel-Hougaard copula*

The covariate term is **W**(*t, β*) = *α*_1_ log(*t*) − *α*_0_ log(*t*) ≡ *β* log(*t*) and so the model is 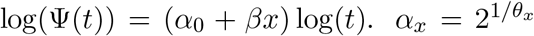, where *θ*_*x*_ ∈ [1, ∞) for *x* = 0, 1 and 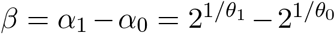. This model indicates that the probability that a candidate is reproducibly ranked among top 100*t*% in workflow *x* = 1 is *t*^*β*^ times of that in *x* = 0. Hence, a negative value of *β* indicates that workflow *x* = 1 is more reproducible than workflow *x* = 0. Here, *c*_1_ is chosen to be 0, 0.1, 0.2, 0.3, 0.4, where a larger *c*_1_ indicates a larger proportion of missing data and *c*_1_ = 0 represents the scenario that all the data are completely observed.

For assessing the type I error, we set *θ*_0_ = *θ*_1_ = 2. For assessing the power, we consider two different settings (*θ*_0_ = 2 and *θ*_1_ = 1.9; *θ*_0_ = 2 and *θ*_1_ = 1.8) to evaluate the performance of the method for different effect sizes. For each case, we simulate 400 datasets, each of which consists of *n* = 10, 000 pairs of observations. We then fit our model with *M* = 30 equally spaced cutoffs in (0, 1) and test *H*_0_ : *β* = 0 using the Wald test at the 5% significance level. Standard errors are calculated using 400 bootstrap samples in case of mdCCR.

As shown in Figure 4, the power of both methods decreases with the increase of the proportion of missing values and the decrease of effect size, when the reproducibility of the two workflows are different. Our method consistently has higher power than CCR for all examined *c*_1_’s. The gain in power is more evident with the increase in the proportion of missing data and the decrease in effect size. The type I errors of mdCCR and CCR are both approximately 0.05 for complete data (*c*_1_ = 1.0 for Nelsen 4.2.12, or *c*_1_ = 0.0 for Gumbel-Hougaard) and then decreases to 0 with the increase in missing data. In addition, the estimates of *α*_0_ and *β* from our method are consistently more accurate with smaller standard errors than those estimated from the CCR approach (Supplementary Tables C7, C8).

**Figure D4:**
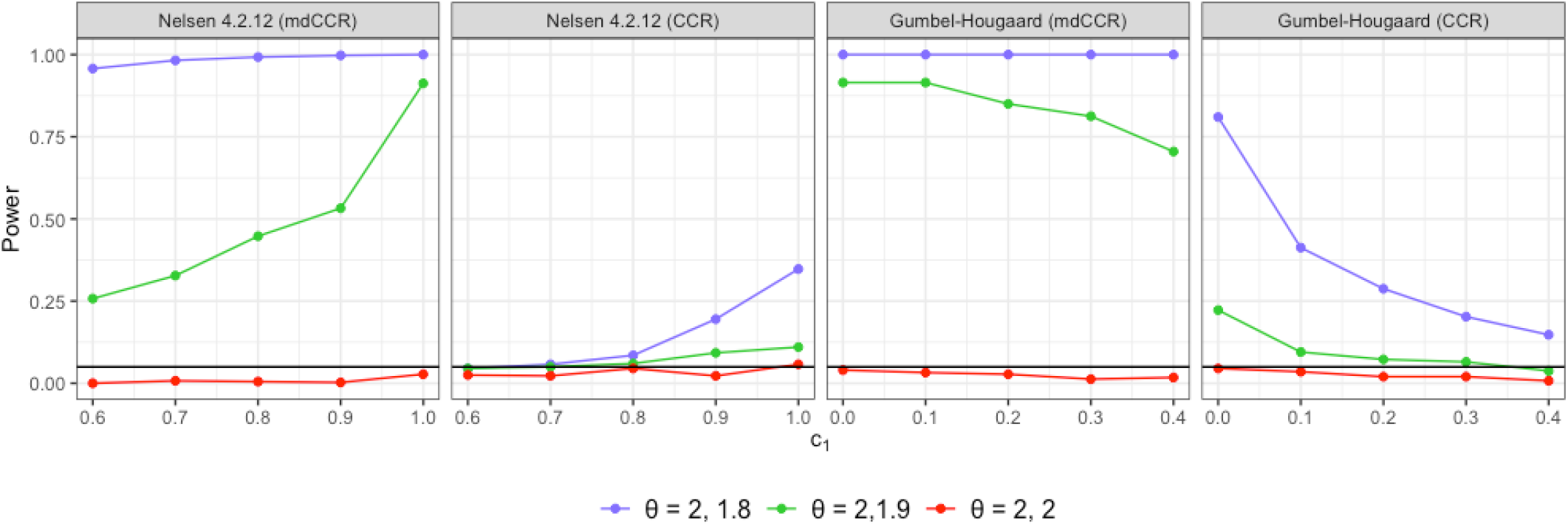
Comparison of power for detecting the difference in reproducibility between two workflows by mdCCR and CCR in simulations. (a, b) are generated from the Nelsen 4.2.12 copula, (c, d) from the Gumbel-Hougaard copula. (a) and (c) are estimated using mdCCR, while (b) and (d) are estimated using the CCR method. Blue (*θ* = 2, 1.8) and green (*θ* = 2, 1.9) lines show the power when the two workflows have different reproducibility. Red line (*θ* = 2, 2) shows the power when the two workflows show no difference in reproducibility, i.e. type I error. The solid black line marks power = 0.05, i.e. the nominal level of type I error. For Nelsen 4.2.12, the amount of missing data increases as *c*_1_ decreases, and for Gumbel-Hougaard, the amount of missing data increases as *c*_1_ increases.

### D.3 Power and accuracy with more than 2 replicates

Next, we assess the power and accuracy for detecting the difference in reproducibility when there are more than two replicates per workflow. Again, two workflows are considered, so *x* takes value 0 or 1 and *θ*_0_ and *θ*_1_ are associated with *x* = 0 and *x* = 1, respectively. We again simulate data from the Gumbel-Hougaard copula and the Nelsen 4.2.12 copula. For each workflow, we simulate R = 1, 3, and 5 pairs of replicates. Because the reproducibility between a pair of replicates from the same workflow may vary across pairs, we incorporate this variation in the simulated data by simulating *θ*_0_ and *θ*_1_ from *N* (2, 0.05) and *N* (1.9, 0.05), respectively. For each case, we simulate 400 datasets, each of which consists of *n* = 10, 000 pairs of observations with varying *c*_1_. We then fit our model (3) with *M* = 30 equally spaced cutoffs in (0, 1) and compute the likelihood to estimate the parameters using (10). We test *H*_0_ : *β* = 0 using the Wald test at the 5% significance level. Standard errors are calculated using 400 bootstrap samples.

As shown in Figure 5, the estimates of *α*_0_ and *β* are more accurate with smaller standard errors when the number of replicates increases (Supplementary Tables C9, C10). Consequently, the power of mdCCR increases with the increase in the number of replicates. The gain in power is more evident with the increase in the proportion of missing data and the number of replicates.

**Figure D5:**
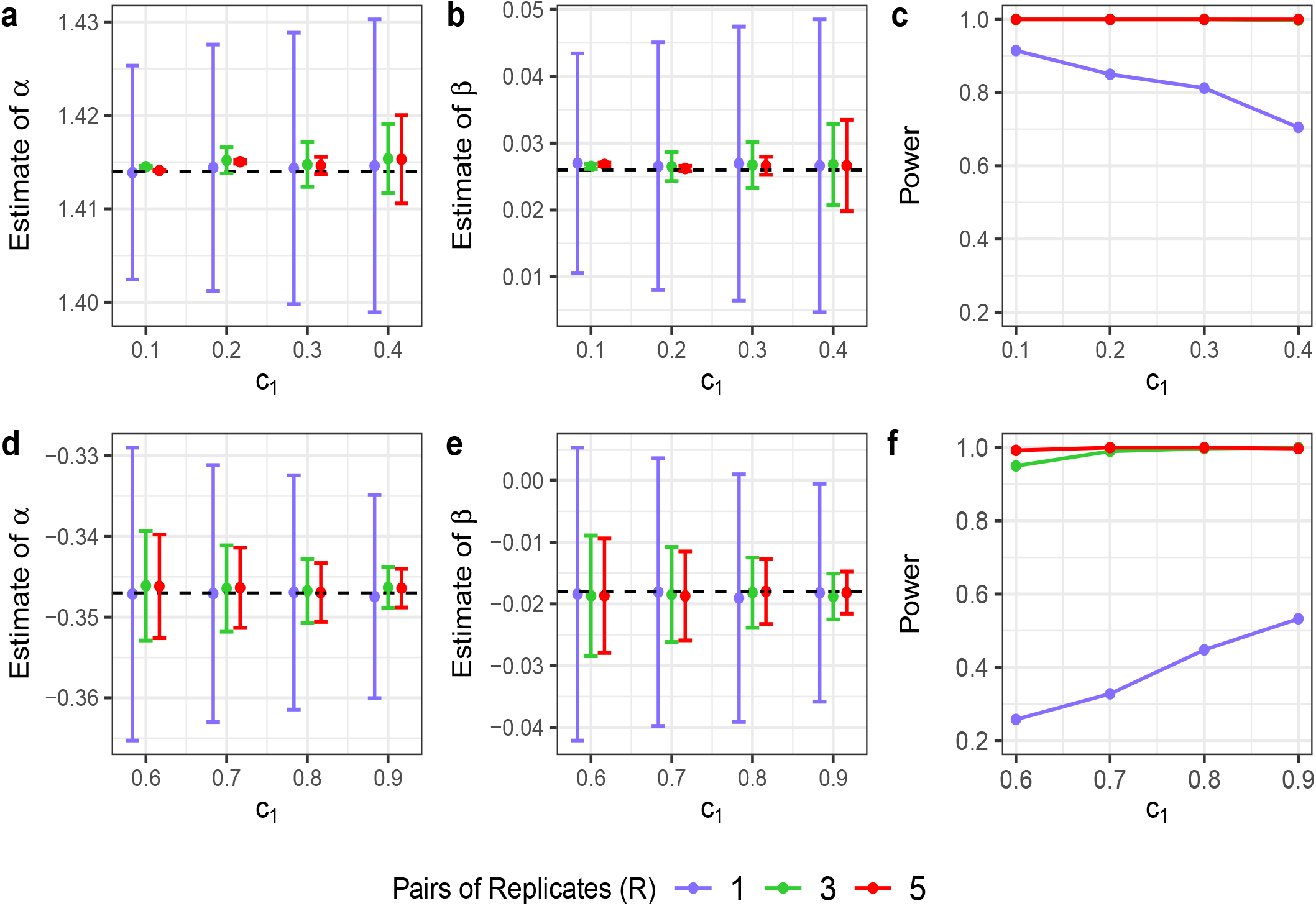
Accuracy in parameter estimation and power of mdCCR for different numbers of replicates. Data are simulated from (a-c) the Gumbel-Hougaard copula and (d-f) the Nelsen 4.2.12 copula, respectively. (a,d) 95% CI of 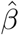; (b,e) 95% CI of 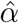; (c,f) power for detecting the difference in reproducibility between workflows. The dashed line in plots (a-b) and (d-e) signifies the true value of the coefficient. For Gumbel-Hougaard, the amount of missing data increases as *c*_1_ increases and for Nelsen 4.2.12, the amount of missing data increases as *c*_1_ decreases. For (c), the power is the same for replicates = 3 and 5, so the green line and the red line overlap in (c).

As previously stated in Section C.4, composite likelihood estimators may exhibit some loss of efficiency when the assumed independence across individual components is violated. We explore this by repeating the simulation above, except that the replicates are reused in different pairs. Specifically, for each workflow, we simulate *R*_0_ = 3 replicates and create *R* = 3 pairs of replicates (i.e. (1,2), (2,3) and (1,3)). These pairs are not entirely independent as each replicate is used in 2 out of 3 pairs. We estimate the parameters and test the significance of *β* in the same way as for the case of independent pairs. Supplementary table C11 shows that the estimates of *α*_0_ and *β* are unbiased for all cases for both copula models, regardless of whether the pairs are dependent or independent. However, it is observed that the variances of 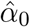 and 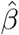 are larger for dependent pairs than independent pairs, leading to some loss in power. This is likely due to the loss of efficiency for composite likelihood estimators when the independence assumption is violated [Varin et al., 2011].

## E Application: Single-cell RNA-seq study of HCT 116 cells

Single-cell RNA-seq (scRNA-seq) is a powerful high-throughput technology recently developed for measuring gene expression profiles of heterogeneous cell populations for each individual cell. It can reveal regulatory relationships between genes and track the trajectories of distinct cell lineages in development at the single-cell resolution [Hwang et al., 2018]. Several experimental factors affect the quality of scRNA-seq data, such as RNA library capture, cell quality and sequencing depth. One of the most important factors in the design of scRNA-seq experiments is to determine the desired sequencing depth [Rizzetto et al., 2017]. If sequencing depth is insufficient, many genes may not be detected. Increasing sequencing depth may improve detection but higher sequencing depth inevitably incurs higher costs [Sims et al., 2014]. Therefore, it is important to find the minimum sequencing depth that allows users to obtain reproducible results. Here, we illustrate the usefulness of our method using a single-cell study of HCT116 cells [Wu et al., 2013].

In this study, the authors generated single-cell transcriptomes from cultured HCT116 cells using different platforms. They made 102 single-cell RNA-seq libraries using 3 methods, including two tube-based methods – (i) *TransPlex Kit (Sigma-Aldrich)* and (ii) *SMARTer Ultra Low RNA Kit (Clontech) for cDNA synthesis*, and one microfluidic method – (iii) *SMARTer cDNA synthesis using the C1 microfluidic system*. They evaluated the sensitivity, reproducibility, and accuracy of these single-cell RNA-seq platforms in comparison with bulk total RNA and multiplexed qPCR. They assessed reproducibility of a platform by computing the proportion of genes detected in different replicates and the Pearson correlation of transcript abundance between different pairs of replicates.

As described in Section A, a major challenge in scRNA-seq data analyses, including the reproducibility assessment, is the large proportion of zeros in the reported gene expression measurements. These zeros occur mainly due to the low amount of mRNA in individual cells and inefficient mRNA capture that are insufficient to produce signals above the detection level, as well as the stochasticity of mRNA expression [Qiu, 2020]. A detailed review on the sources of zeros in single-cell RNA-seq data can be found in [Jiang et al., 2022]. Though several statistical methods have been developed for imputing zeros in scRNA-seq data, a recent study [Sarkar and Stephens, 2021] pointed out that imputing zeros based on the missing mechanisms, such as, missing completely at random or missing at random, should be avoided, because these missing mechanisms, which imply gene expression values are randomly recorded as zeros regardless the underlying true expression level, do not happen in the scRNA-seq measurement process. Our model nevertheless provides a way to take account of the zeros in the reproducibility assessment without imputation. We will use our method to investigate the effect of platforms and sequencing depth on reproducibility and compare with CCR.

### E.1 Comparing the reproducibility of tube-based and microfluidic-based libraries

For simplicity, we will refer to method (i) as TransPlex, (ii) as SMARTer, and (iii) as C1. We consider these 3 experimental platforms as different workflows and compare their reproducibility between cells using our method. In this study, C1 has 96 single cells, but TransPlex and SMARTer have only 3 single cells for each platform. Therefore, we formed 3 pairs of replicates for each workflow using 3 single cells for TransPlex, 3 for SMARTer and 6 for C1. On average, each replicate for SMARTer and TransPlex has approximately 50 million raw reads while that for C1 has approximately 3 million raw reads. The average number of zeros in each pair of replicates from the 3 workflows (TransPlex, SMARTer and C1) are 11,794, 14,022 and 15,119, respectively. For CCR, we only used a single pair of single cells for each platform as the CCR setup does not include the case for more than 2 replicates. We examine the rank scatterplots of the scores on the two biological replicates to select the functional form of the regression model and find that the Gumbel-Hougaard copula is a reasonable fit (see Supplementary Figure B7). Therefore, we fit the following regression model,

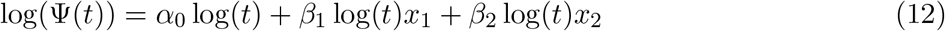

using *M* = 30 equally spaced cutoffs in (0,1), where TransPlex is used as the baseline and *x*_1_, *x*_2_ are dummy variables for SMARTer and C1, respectively.

Table E4 summarizes the results from our analysis. According to the 95% confidence intervals of *β*’s for mdCCR, the three workflows are significantly different in reproducibility. Among them, C1 is the most reproducible and SMARTer is the least reproducible. For example, at *t* = 0.05, the estimated probabilities of being reproducibly reported in the top 100*t*% are *t*^−0.074^ = 1.25 and *t*^−0.068^ = 1.23 times higher for C1 and SMARTer as for TransPlex, respectively. This agrees with the results in Wu et al. (2013) [Wu et al., 2013], whose assessment also indicated a high proportion of reproducibly detected genes and a high correlation between different pairs of replicates generated by the C1 platform. However, as discussed in Section A, different types of correlation (with or without zeros) can lead to different conclusions depending upon the choice of pairs of replicates. Nevertheless, our approach uses a principled way to take missing data and all replicates into account and provides quantitative information on the reproducibility at any given threshold. While the estimates of CCR show a similar trend as those of mdCCR, they fail to detect a statistically significant difference in reproducibility between SMARTer and Transplex at the 95% confidence level. This lack of power in CCR is likely related to its ignorance of missing data and restriction to the concordance of a single pair of replicates, as observed in the simulation studies (Section D).

**Table E4:** Estimated effect of different libraries by mdCCR and CCR, using Transplex as the baseline.

|  |  | mdCCR |  |  |  | CCR |  |
| --- | --- | --- | --- | --- | --- | --- | --- |
| Platform | | Estimate | 95% CI | $\hat{c}_1$ | $\hat{c}_2$ | Estimate | 95% CI |
| TransPlex | $\alpha_0$ | 1.375 | (1.353, 1.397) | 0.648 | 0.534 | 1.531 | (1.500, 1.563) |
| SMARTer | $\beta_1$ | -0.068 | (-0.086, -0.050) | 0.678 | 0.585 | -0.020 | (-0.070, 0.029) |
| C1 | $\beta_2$ | -0.074 | (-0.103, -0.045) | 0.663 | 0.663 | -0.049 | (-0.097, -0.001) |

### E.2 Finding the required sequencing depth to achieve sufficient reproducibility

Next, we investigate the effect of sequencing depth on reproducibility. In particular, we aim to find the minimum sequencing depth required to obtain reproducible results. Because Section E.1 shows that the C1 microfluidic system is the most reproducible platform among the three platforms, we used the C1 data for this analysis. We took 10 cells from C1 data to form 5 pairs of replicates. Their sequencing depth ranges from 1.6M to 5M.

Because differences in sequencing depth across replicates can confound with the reproducibility analysis, we first downsample all the replicates to a common depth of 1.6M to ensure equality across replicates. To study how sequencing depth affects the reproducibility across replicates, we downsample the reads in the data to create a series of sequencing depths, a common way to study the effect of sequencing depth. Here we downsample the data from 1.6M reads to .8M, 1M, 1.2M, 1.4M reads, and compare the reproducibility across each sequencing depth using our method. See the Appendix for details on the data processing procedure. The number of zeros increases with the reduction of sequencing depth, ranging from 14,000 to 17,000. We consider different downsampling levels as different workflows. Rank scatterplots show that the Gumbel-Hougaard copula fits each downsampled dataset well. Using *M* = 30 equally spaced cutoffs in (0,1), we fit the following model,

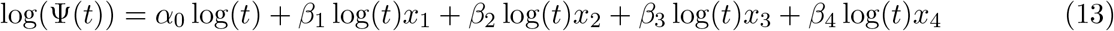

where the original data (1.6M) is used as the baseline and *x*_1_, *x*_2_, *x*_3_, and *x*_4_ are dummy variables for 1.4M, 1.2M, 1M and 0.8M depth, respectively.

The estimated parameters are summarized in Table E5 and Figure 6. According to the 95% confidence intervals of *β*’s for mdCCR, we observe that when the sequencing depth is below 1M, the workflows are significantly less reproducible than the original depth (1.6M). However, after the sequencing depth reaches 1M, the reduction in reproducibility from the original depth is no longer significant. This indicates that a minimal sequencing depth of 1M is required to reach the reproducibility level that is not significantly lower than the original sample of 1.6M reads.

**Table E5:** Estimated effect of different sequencing depth by mdCCR, using 1.6M as the baseline.

| Sequencing Depth | Coefficient | Estimate | 95% CI | $\hat{c}_1$ | $\hat{c}_2$ |
| --- | --- | --- | --- | --- | --- |
| Original (1.6M) | $\alpha_0$ | 1.3032 | (1.2891, 1.3173) | 0.677 | 0.663 |
| 1.4M | $\beta_1$ | 0.0026 | (-0.0137, 0.0189) | 0.683 | 0.668 |
| 1.2M | $\beta_2$ | 0.0072 | (-0.0089, 0.0233) | 0.694 | 0.679 |
| 1.0M | $\beta_3$ | 0.0095 | (-0.0066, 0.0256) | 0.699 | 0.685 |
| 0.8M | $\beta_4$ | 0.0161 | (0.0002, 0.0320) | 0.704 | 0.689 |

**Figure E6:**
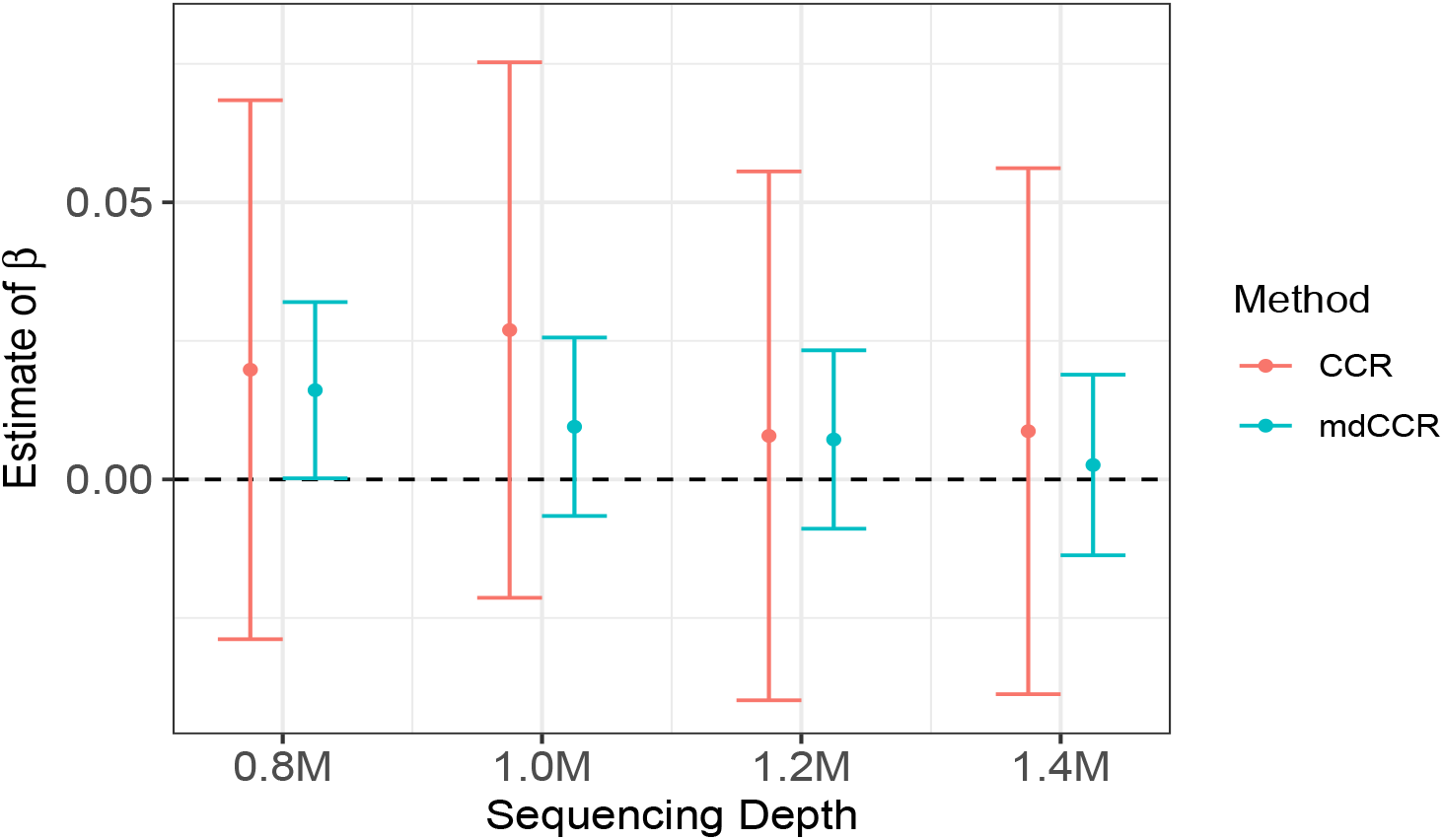
Evaluation of sequencing depth effect on reproducibility for HCT116 single cell RNA-seq data generated using the C1 platform: 95% confidence interval for 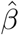 coefficients at different sequencing depths. The results from one pair of replicates (pair 1 in the Supplementary Table C12) are plotted for CCR.

In contrast, unlike mdCCR, which analyzes all replicates at once and provide a unified result, CCR gives different results for different pairs of replicates. Results from CCR using 2 different pairs of replicates are provided in the Supplementary Table C12. It is evident that CCR does not detect the lack of sufficiency when sequencing depth is below 1M.

## F Conclusion

In this paper, we present a regression framework for assessing the influence of experimental factors on the reproducibility of high-throughput experiments when there are missing data. By using a principled approach to take account of missing values, the lack of reproducibility reflected by the missing values is properly accounted for in the assessment of the influence of experimental factors. Using our method, we compared the reproducibility of different single-cell platforms and identified the minimum sequencing depth that can generate sufficiently reproducible results for a scRNA-seq sample. This shows that our method can be used as a principled general approach to understand how influencing factors affect the reproducibility of high-throughput experiments.

Our approach can also be applicable to other experiments where missing values are prevalent, such as Functional Magnetic Resonance Imaging (fMRI) data. The fMRI measures brain activity by detecting changes associated with blood flow using the blood-oxygen-level-dependent (BOLD) contrast. The BOLD signal is frequently corrupted by noise from multiple artifacts such as imaging hardware and temperature, leading to weak or absent signals [Yan et al., 2020, Sammartino et al., 2016]. This hampers studies that investigate whole-brain activity and may result in misinterpretation of findings. The importance of operational factors on the reproducibility of a BOLD signal can be evaluated using our proposed method to help construct reliable data generation protocols.

## Acknowledgments

This article is supported by NIH R01GM109453.

## Data Availability

The raw data can be obtained from Gene Expression Omnibus using accession code GSE51254.

## A Supplementary Information

### A.1 Estimation Procedure: Maximum likelihood estimation via the EM algorithm

#### EM algorithm for one pair of replicates from a single workflow

First, we consider the case of missing data framework for a single workflow and a single pair of replicates. We first consider the framework in Section C.1.

As discussed in Sections C.1 and C.2, the sample space can be partitioned into categories *C*11, *C*12, *C*21 and *C*22, where the observations in *C*11 are completely observed, those in *C*12 and *C*21 are partially observed, and those in *C*22 are completely missing. The numbers of occurrence in these categories, *n*_11_, *n*_12_ and *n*_21_, are observed, but *n*_22_ is missing. The category *C*11 can be partitioned into *C*11^*a*^ and *C*11^*b*^, where *C*11^*a*^ can be further partitioned into *M* blocks (*C*11*m, m* = 1, 2, …, *M*) by the cutoffs, 0 < *t*_1_ < …< *t*_*M*_ = *c*_1_. The corresponding probabilities for the categories *C*11^*a*^, *C*11^*b*^, *C*12, and *C*21 are

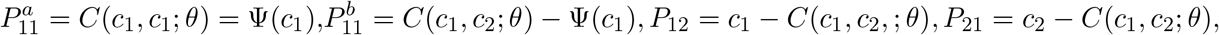

respectively, and that for *C*11*m* is *P*_11*m*_(*ϕ*) = Ψ(*t*_*m*_) − Ψ(*t*_*m*−1_), where Ψ(·) = *g*^−1^(Σ*α*_*k*_*h*_*k*_(·) + **W**(·, ***β***)′**x**) and *C*(·, ·; *θ*) is the copula function corresponding to the regression model, with the copula association parameter *θ* obtained from its relation with *α* and *β* (See Section B.1). For example, the Nelsen 4.2.12 copula fits this framework and its copula function is defined as 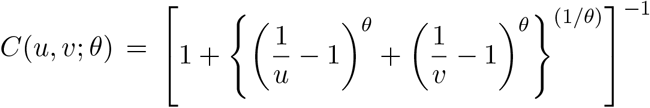, where the association parameter for a single workflow is 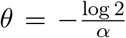 (see Table B2). In the special case of *c*_1_ = *c*_2_, the above probabilities can be computed directly from Ψ(·) from the regression model without using the copula function *C*(·, ·; *θ*).

Denote the parameters to be estimated as *ϕ* = {***ω***, *c*_1_, *c*_2_}, where ***ω*** = (***α, β***). We estimate them using maximum likelihood approach for truncated data via the EM algorithm [Dempster et al., 1977]. Denote the number of observed data from a workflow as *n*_*obs*_ = *n*_11_ + *n*_12_ + *n*_21_ and *P*(*ϕ*) = *P*_11_(*ϕ*) + *P*_12_(*ϕ*) + *P*_21_(*ϕ*) = *c*_1_ + *c*_2_ − *C*(*c*_1_, *c*_2_; *θ*), then the observed data from a workflow can be represented as 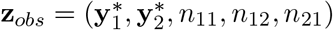, where 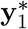 and 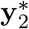 are observed scores on replicates 1 and 2, respectively.

Following the EM algorithm for truncated data [Dempster et al., 1977], the number of unobserved candidates in category *C*22, *n*_22_ ∼ Negative Binomial(*n*_*obs*_, *P*(*ϕ*)), i.e.,

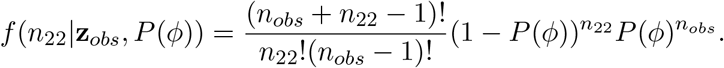

The observed data likelihood is,

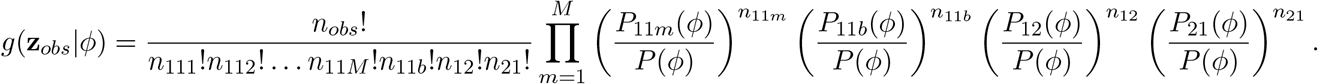

Here 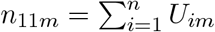, where *U*_*im*_ = *I*(*i* ∈ 𝒴_*m*_|**x**, *c*_1_, *c*_2_) and 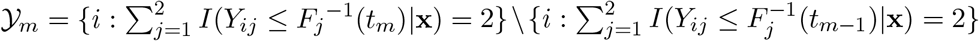. Because *F*_*j*_(·) is not observable due to truncation, it is computed as 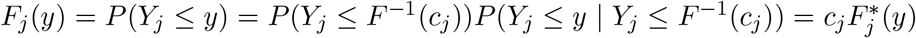 for *y* ≤ *F*^−1^(*c*_*j*_), where 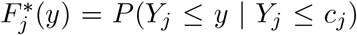 is cumulative distribution of the truncated distribution and can be computed from the observed scores. (Here we use the empirical distribution of 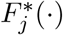 in our implementation.) Hence, *n*_11*m*_ is a function of *t*_*m*_, *c*_1_ and *c*_2_, and needs to be updated when the estimates of *c*_1_ and *c*_2_ are updated, unlike the case when all data points are completely observed, where *n*_11*m*_ is known and fixed given *t*_*m*_. Similarly, *n*_11*b*_ is also dependent on *c*_1_ and *c*_2_, and needs to be updated when the estimates of *c*_1_ and *c*_2_ are updated.

We augment the data to include the missing counts, *n*_22_, then the complete data is **z**_*c*_ = {**z**_*obs*_, *n*_22_}. The complete data likelihood is,

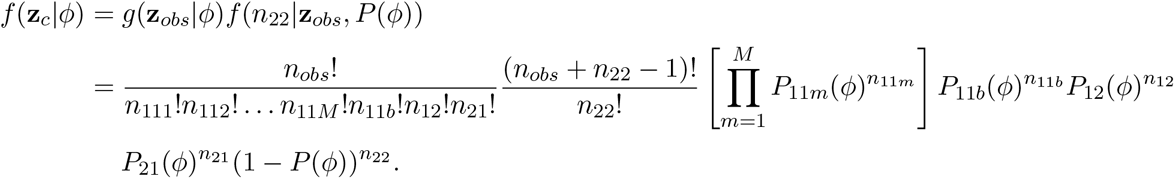

Following the EM methodology for truncation [Dempster et al., 1977], the M-step optimizes the function, *Q*(*ϕ*|*ϕ*^(*p*)^) = E[log *f*(**z**_*c*_|*ϕ*)|**z**_*obs*_, *ϕ*^(*p*)^], where

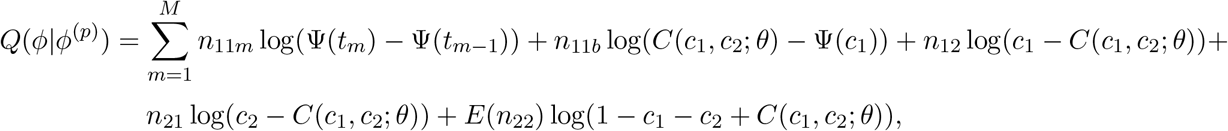

and the E-step is defined as,

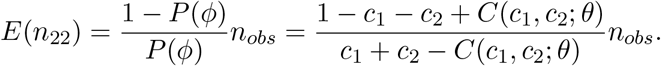

For the inverted missing data framework in Section C.3, the probabilities of falling in category *C*11*m* are *P*_11*m*_(*ϕ*) = Ψ*(*t*_*m*−1_) − Ψ*(*t*_*m*_); *m* = 1, 2, …, *M*, where Ψ*(*t*_*m*_) = 1 − 2*t*_*m*_ + Ψ(*t*_*m*_). The probability of falling in category *C*11^*b*^, *C*12 and *C*21 is *P*_11*b*_ = *c*_1_ − *c*_2_ + *C*(*c*_1_, *c*_2_; *θ*) − Ψ(*c*_1_), *P*_12_(*ϕ*) = *c*_2_ − *C*(*c*_1_, *c*_2_; *θ*), and *P*_21_(*ϕ*) = *c*_1_ − *C*(*c*_1_, *c*_2_; *θ*), respectively, where *C*(·, ·; *θ*) is the copula function corresponding to the regression model, with the association parameter *θ* obtained from its relation with *α* and *β*. For example, the Gumbel-Hougaard copula fits this framework and its copula function is defined as 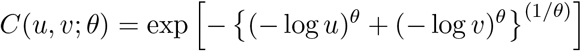, where the association parameter for a single workflow is 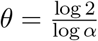 (See Table B2). Now, the function for M-step is defined as,

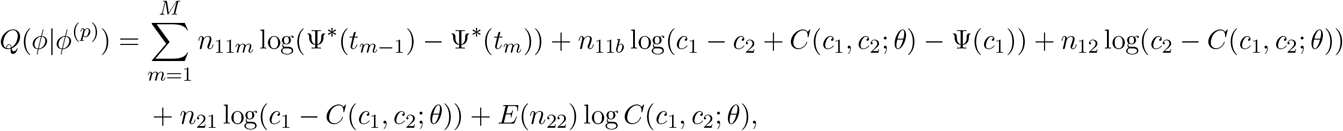

and the E-step is defined as,

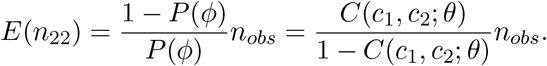

#### EM algorithm for *S* workflows each with one pair of replicates

Now, let us consider the case of *S* workflows. The number of observed data from *S* workflows is denoted as 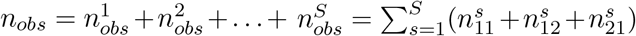 and the number of candidates in category *C*22 of the *s*^*th*^ workflow, 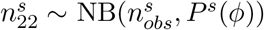. Here, the parameters to be estimated are 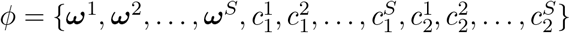. So, the M-step optimizes the following function,

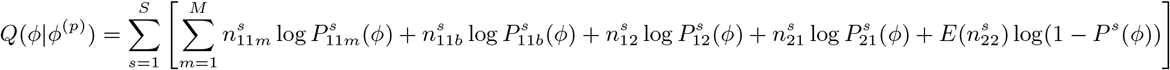

and the E-step is defined as,

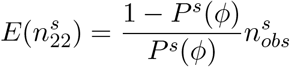

where 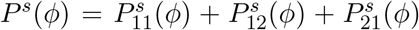 and 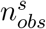 is the total number of candidates observed from workflow *s*.

#### EM algorithm for *S* workflows each with *R* pairs of replicates

When each workflow has *R* pairs of replicates, the number of observed data from *S* workflows is denoted as 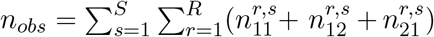. For a given workflow *s*, we assume that all pairs of replicates share the same set of reproducibility parameters, ***ω***^*s*^, and truncation parameter, 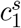, because all replicates are generated from the same workflow. Hence, the number of unobserved candidates for *s*^*th*^ workflow from all replicates share the same distribution, i.e., 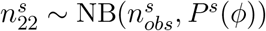. Consequently, the number of parameters to be estimated remain the same i.e., 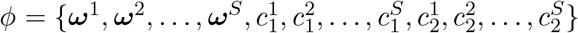, but these parameters are estimated based on the observations from all the R pairs of replicates. Specifically, the function to be optimized in the M-step is

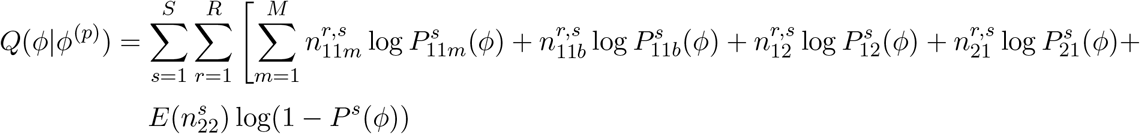

and the E-step is defined as

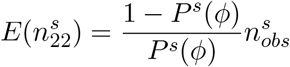

where 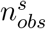 is the average number of candidates observed from workflow *s* across *R* pairs of replicates and 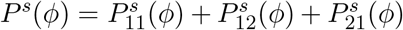. Algorithm 1 describes the general algorithm to estimate *ϕ* for *S* workflows with *R* pairs of replicates. We use bootstrapping method for calculating the standard errors.

##### Algorithm 1 EM algorithm for estimating *ϕ*

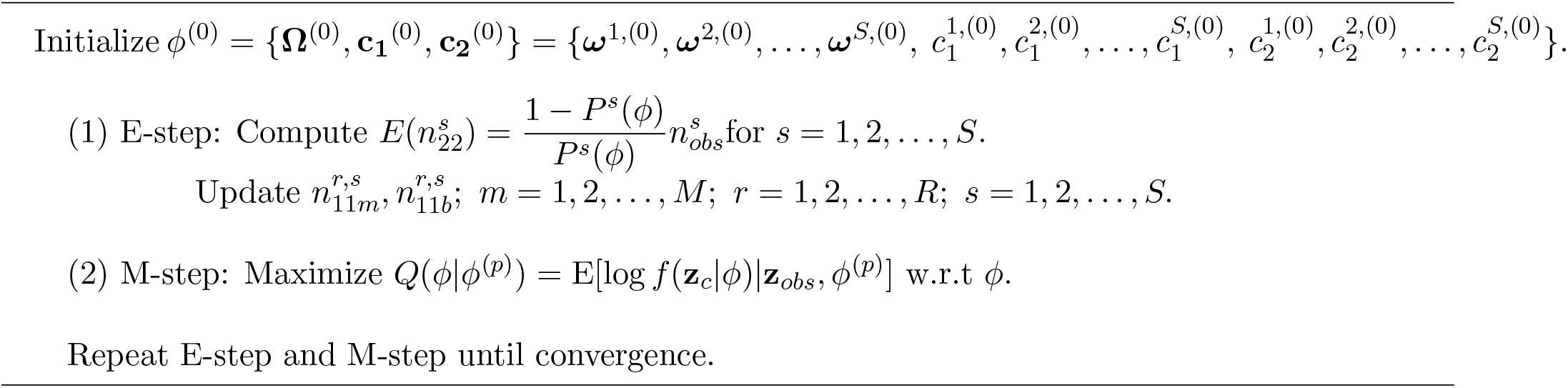

### A.2 Downsampling and pre-processing of raw reads

The raw data were obtained from Gene Expression Omnibus using accession code GSE51254. We downsampled the data (randomly selected raw reads from the data) using Seqtk [Li, 2012] and then removed the adapter sequences using Trim Galore [Krueger, 2015]. Alignment and mapping were done using Bowtie 2 [Langmead and Salzberg, 2012] against the reference genome, hg19. Gene expression was measured using featureCounts [Liao et al., 2013]. All the pre-processing steps were done using usegalaxy.org [Afgan et al., 2018].

## B Supplementary Figures

**Figure B7:**
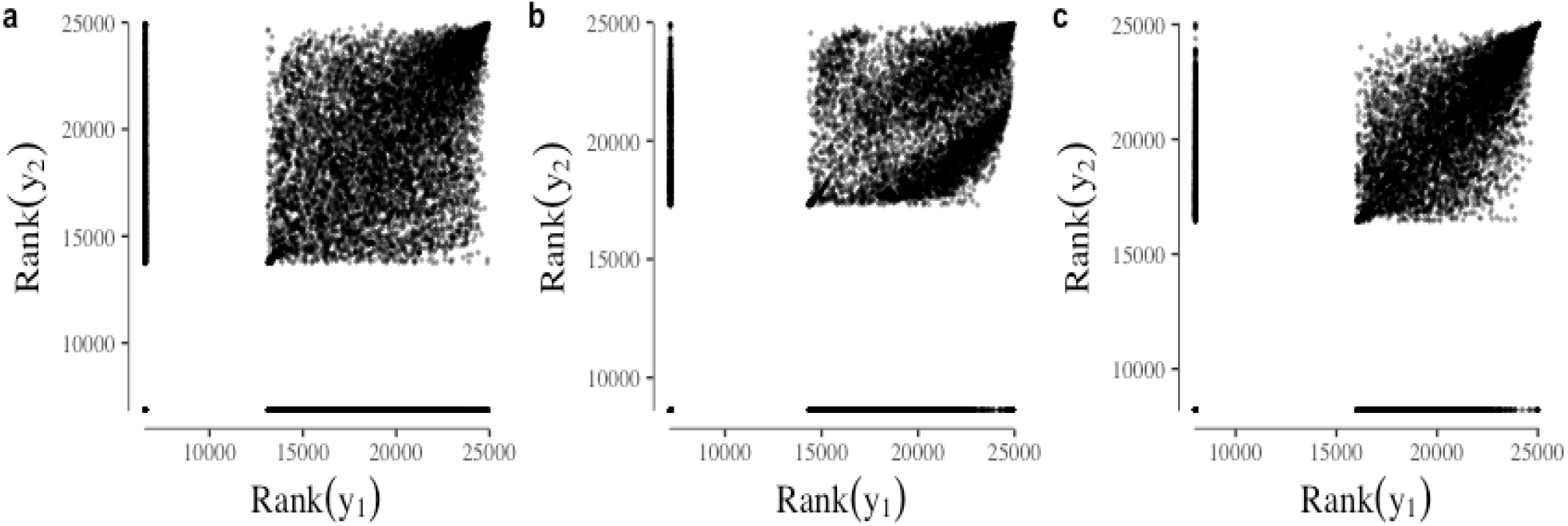
Rank scatterplots of gene expression on two replicates from (a) TransPlex, (b) SMARTer and (c) C1. The dots on the left and bottom sides of the plot (appearing like lines) are the partial data points as the genes are only observed on a single replicate. This resembles the inverted framework described in Section 3.3.

## C Supplementary Tables

**Table C6:** Proportion of zeros and correlation coefficients calculated for 2 replicates from 3 platforms. Correlation coefficients are measured with and without the zeros.

| Platform | Proportion of zeros | Spearman's rank correlation |  | Pearson's correlation |  |
| --- | --- | --- | --- | --- | --- |
|  |  | With 0 | Without 0 | With 0 | Without 0 |
| TransPlex | 0.430 | 0.648 | 0.501 | 0.962 | 0.984 |
| SMARTer | 0.612 | 0.734 | 0.460 | 0.477 | 0.789 |
| C1 | 0.568 | 0.766 | 0.751 | 0.685 | 0.984 |

**Table C7:**
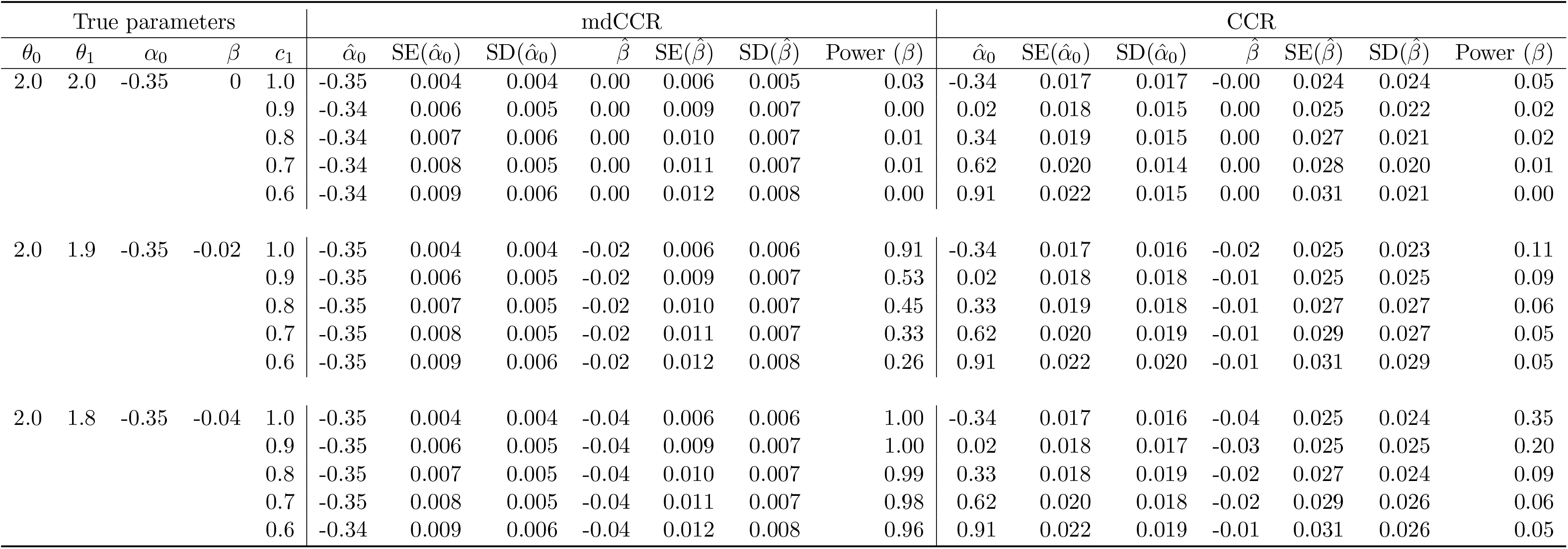
Estimation of *α* and *β* in the simulation study of two workflows for different values of *c*_1_ in the Nelsen 4.2.12 copula, using our proposed method (mdCCR) and CCR method.

**Table C8:**
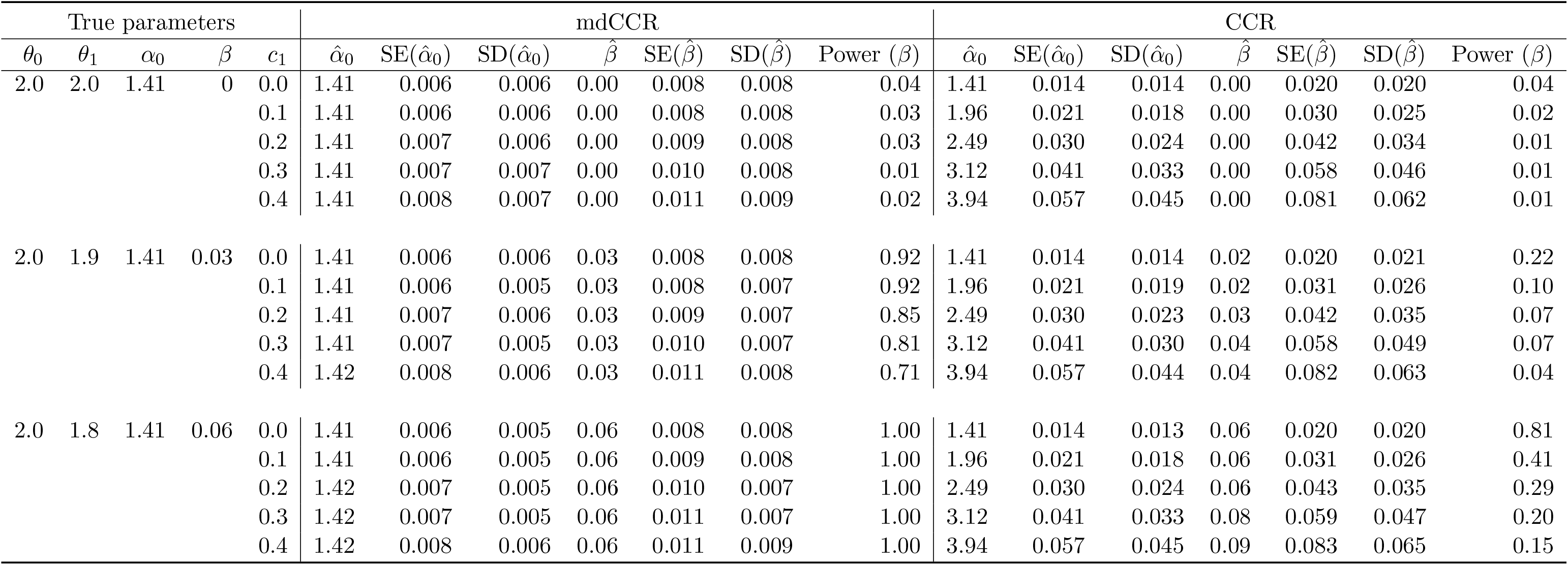
Estimation of *α* and *β* in the simulation study of two workflows for different values of *c*_1_ in the Gumbel-Hougaard copula, using our proposed method (mdCCR) and CCR method.

**Table C9:** Estimation of *α* and *β* in the simulation study of two workflows with *R* = 1, 3 and 5 pairs of replicates per workflow for different values of *c*_1_ in the Nelsen 4.2.12 copula, using the mdCCR method. Here, *θ*_0_ = 2 and *θ*_1_ = 1.9 i.e. *α*_0_ = −0.347 and *β* = −0.018.

| R | $c_1$ | $\hat{\alpha}_0$ | SE($\hat{\alpha}_0$) | SD($\hat{\alpha}_0$) | $\hat{\beta}$ | SE($\hat{\beta}$) | SD($\hat{\beta}$) | Power ($\beta$) |
| --- | --- | --- | --- | --- | --- | --- | --- | --- |
| 1 | 0.9 | -0.347 | 0.006 | 0.005 | -0.018 | 0.009 | 0.007 | 0.53 |
|  | 0.8 | -0.346 | 0.007 | 0.005 | -0.018 | 0.010 | 0.007 | 0.45 |
|  | 0.7 | -0.346 | 0.008 | 0.005 | -0.019 | 0.011 | 0.007 | 0.33 |
|  | 0.6 | -0.346 | 0.009 | 0.006 | -0.019 | 0.012 | 0.008 | 0.26 |
| 3 | 0.9 | -0.346 | 0.001 | 0.003 | -0.019 | 0.002 | 0.005 | 1.00 |
|  | 0.8 | -0.347 | 0.002 | 0.003 | -0.018 | 0.003 | 0.004 | 0.99 |
|  | 0.7 | -0.346 | 0.003 | 0.003 | -0.018 | 0.004 | 0.005 | 0.99 |
|  | 0.6 | -0.346 | 0.003 | 0.003 | -0.019 | 0.005 | 0.005 | 0.95 |
| 5 | 0.9 | -0.346 | 0.001 | 0.003 | -0.018 | 0.002 | 0.004 | 1.00 |
|  | 0.8 | -0.347 | 0.002 | 0.003 | -0.018 | 0.003 | 0.004 | 1.00 |
|  | 0.7 | -0.346 | 0.003 | 0.003 | -0.019 | 0.004 | 0.004 | 1.00 |
|  | 0.6 | -0.346 | 0.003 | 0.003 | -0.019 | 0.005 | 0.004 | 0.99 |

**Table C10:**
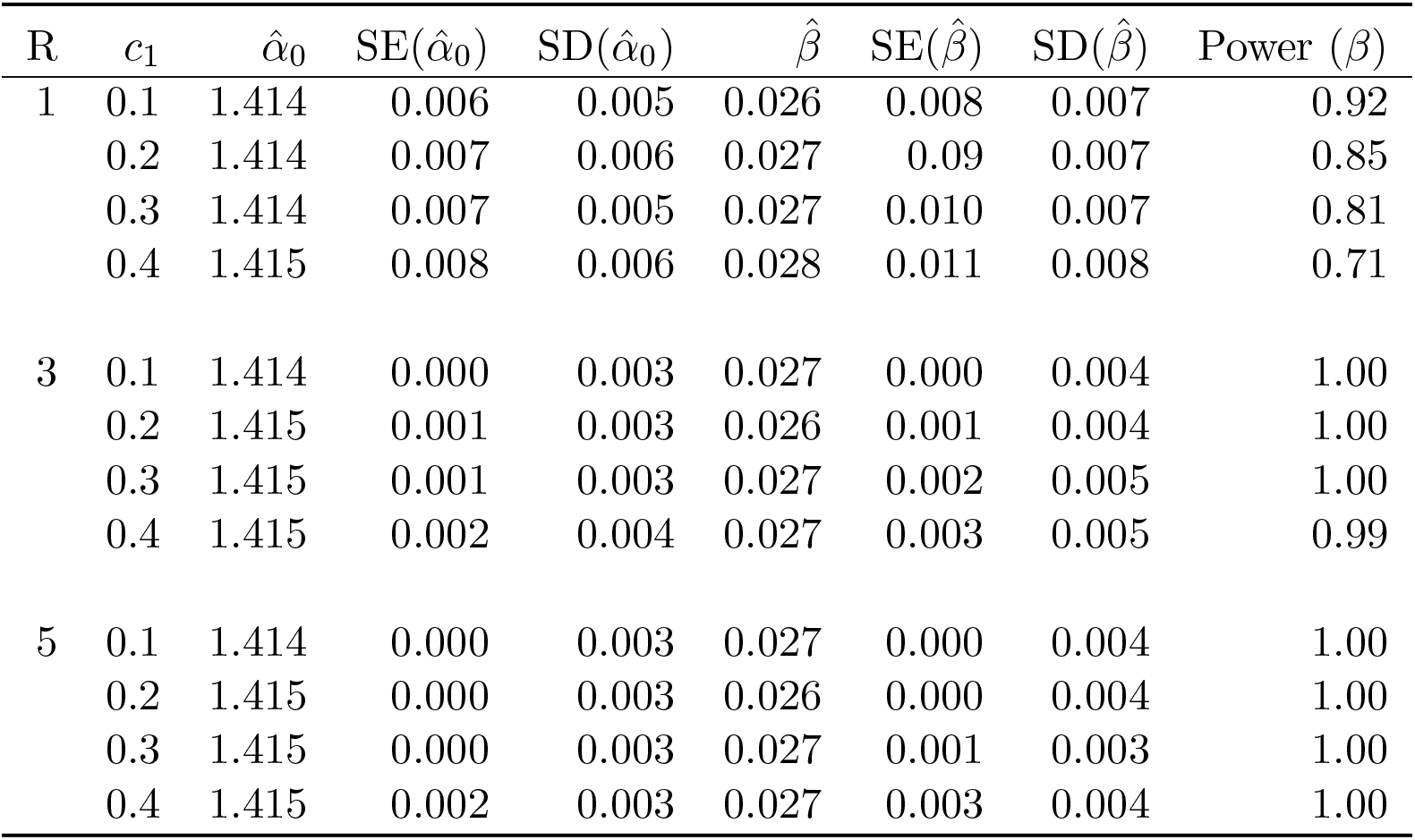
Estimation of *α* and *β* in the simulation study of two workflows with *R* = 1, 3 and 5 pairs of replicates per workflow for different values of *c*_1_ in the Gumbel-Hougaard copula, using the mdCCR method. Here, *θ*_0_ = 2 and *θ*_1_ = 1.9 i.e. *α*_0_ = 1.414 and *β* = 0.026.

**Table C11:**
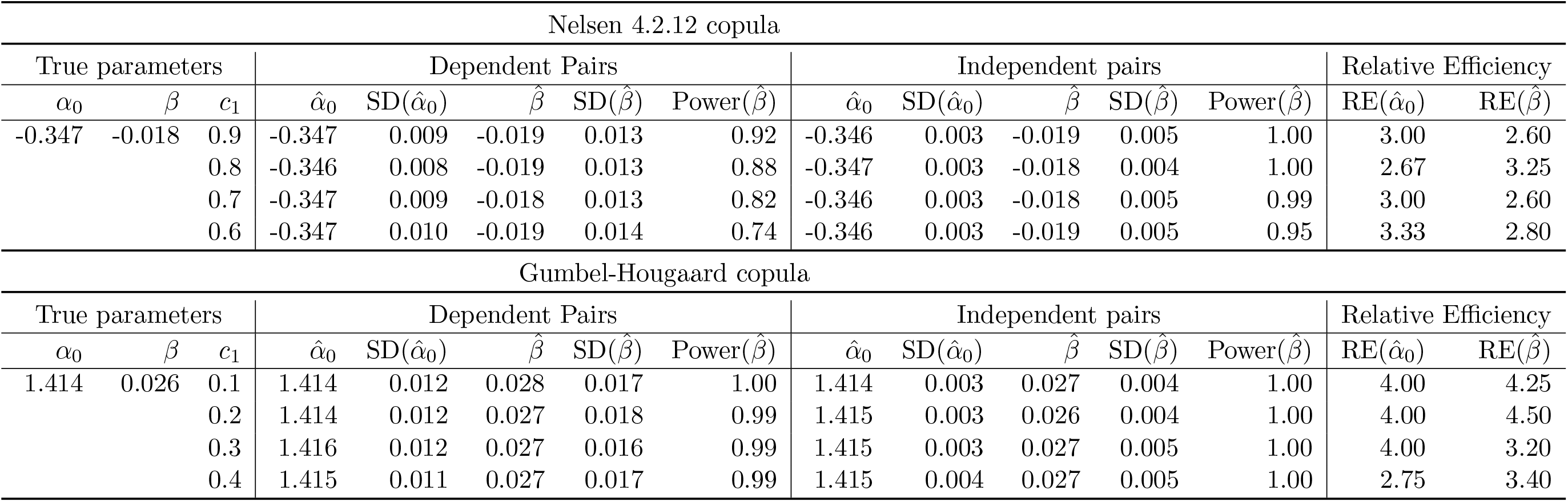
Estimation of *α* and *β* in the simulation study of two workflows with *R* = 3 independent (created using 6 replicates) and dependent (created using 3 replicates only) pairs of replicates per workflow for different values of *c*_1_ in the Nelsen 4.2.12 and Gumbel-Hougaard copula, using the mdCCR method. Here, *θ*_0_ = 2 and *θ*_1_ = 1.9. Relative efficiency 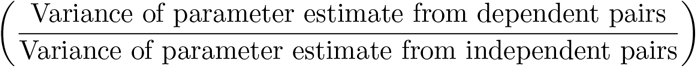 is reported.

**Table C12:** Estimated effect of different sequencing depth by CCR, using 1.6M as the baseline. Table shows two different results based on two different pairs of replicates.

|  | Sequencing Depth | Coefficient | Estimate | 95% CI |
| --- | --- | --- | --- | --- |
| Pair 1 | Original (1.6M) | $\alpha_0$ | 1.410 | (1.380,1.447) |
| | 1.4M | $\beta_1$ | 0.009 | (-0.039,0.056) |
| | 1.2M | $\beta_2$ | 0.008 | (-0.040,0.056) |
| | 1.0M | $\beta_3$ | 0.027 | (-0.021, 0.075) |
| | 0.8M | $\beta_4$ | 0.020 | (-0.029, 0.068) |
| Pair 2 | Original (1.6M) | $\alpha_0$ | 1.500 | (1.465,1.537) |
| | 1.4M | $\beta_1$ | -0.003 | (-0.054,0.049) |
| | 1.2M | $\beta_2$ | 0.003 | (-0.049,0.055) |
| | 1.0M | $\beta_3$ | -0.005 | (-0.057, 0.047) |
| | 0.8M | $\beta_4$ | -0.017 | (-0.070, 0.035) |

